# ATP sensing in living plant cells reveals tissue gradients and stress dynamics of energy physiology

**DOI:** 10.1101/153163

**Authors:** Valentina De Col, Philippe Fuchs, Thomas Nietzel, Marlene Elsässer, Chia Pao Voon, Alessia Candeo, Ingo Seeliger, Mark D. Fricker, Christopher Grefen, Ian Max Møller, Andrea Bassi, Boon Leong Lim, Marco Zancani, Andreas J. Meyer, Alex Costa, Stephan Wagner, Markus Schwarzländer

**Author notes:** **Corresponding authors:** Stephan Wagner; Ph. Markus Schwarzländer; Ph.

## Abstract

Growth and development of plants is ultimately driven by light energy captured through photosynthesis. ATP acts as universal cellular energy cofactor fuelling all life processes, including gene expression, metabolism, and transport. Despite a mechanistic understanding of ATP biochemistry, ATP dynamics in the living plant have been largely elusive. Here we establish live MgATP^2−^ assessment in plants using the fluorescent protein biosensor ATeam1.03-nD/nA. We generate Arabidopsis sensor lines and investigate the sensor *in vitro* under conditions appropriate for the plant cytosol. We establish an assay for ATP fluxes in isolated mitochondria, and demonstrate that the sensor responds rapidly and reliably to MgATP^2−^ changes *in planta*. A MgATP^2−^ map of the Arabidopsis seedling highlights different MgATP^2−^ concentrations between tissues and in individual cell types, such as root hairs. Progression of hypoxia reveals substantial plasticity of ATP homeostasis in seedlings, demonstrating that ATP dynamics can be monitored in the living plant.

**One-sentence Summary:** Sensing of MgATP^2−^ by fluorimetry and microscopy allows dissection of ATP fluxes of isolated organelles, and dynamics of cytosolic MgATP^2−^ *in vivo*.

**Funding Agencies:** This work was supported by the Deutsche Forschungsgemeinschaft (DFG) through the Emmy-Noether programme (SCHW1719/1-1; M.S. and GR4251/1-1; C.G.), the Research Training Group GRK 2064 (M.S.; A.J.M.), the Priority Program SPP1710 (A.J.M.) and a grant (SCHW1719/5-1; M.S.) as part of the package PAK918. The Seed Fund grant CoSens from the Bioeconomy Science Center, NRW (A.J.M.; M.S.) is gratefully acknowledged. The scientific activities of the Bioeconomy Science Center were financially supported by the Ministry of Innovation, Science and Research within the framework of the NRW Strategieprojekt BioSC (No. 313/323-400-002 13). A.Co. received funding by the Ministero dell’Istruzione, dell’Università e della Ricerca through the FIRB 2010 programme (RBFR10S1LJ_001) and Piano di Sviluppo di Ateneo 2015 (Università degli Studi di Milano). M.Z. received funding by the Ministero dell’Istruzione, dell’Università e della Ricerca (Italy) through the PRIN 2010 programme (PRIN2010CSJX4F). S.W. and T.N. received travel support by the Deutscher Akademischer Austauschdienst (DAAD). V.D.C. was supported by the European Social Fund, Operational Programme 2007/2013, and an Erasmus+ Traineeship grant. M.D.F was supported by The Human Frontier Science Program (RPG0053/2012), and the Leverhulme Foundation (RPG-2015-437). I.M.M. was supported by a grant from the Danish Council for Independent Research - Natural Sciences. V.C.P. was supported by the Innovation and Technology Fund (Funding Support to Partner State Key Laboratories in Hong Kong) of the HKSAR.

**Abbreviations:** AAC – ADP/ATP carrier; AK – adenylate kinase; cAT – carboxyatractyloside; CCCP – carbonyl cyanide m-chlorophenyl hydrazone; CFP – cyan fluorescent protein; CLSM – confocal laser scanning microscopy; ETC – electron transport chain; FRET – Förster Resonance Energy Transfer; LSFM – light sheet fluorescence microscopy.

## Introduction

ATP is universal in cells. It is used as a metabolic building block and as a cofactor to couple exergonic and endergonic reactions, making ATP as fundamental to life as proton gradients and the genetic code. An additional function as a biological hydrotrope to keep proteins soluble was recently suggested (Patel et al., 2017). De-phosphorylation to ADP and AMP, and re-phosphorylation to ATP allow high energy fluxes based on relatively small pool sizes in the cell (Rich, 2003). The major sites of ATP synthesis are ATP synthases driven by the proton motive force established across the inner membrane of the mitochondria, and the plastid thylakoids in plants. The ATP produced typically fuels the wide range of energy-demanding processes, such as motility, transport, and gene expression, in other parts of the cell, although plastids also consume substantial amounts of ATP in the Calvin-Benson cycle for example, to an extent that varies substantially with the light/dark cycle. Coupling of these interactions requires ATP/ADP exchange between the organelles and cytosol. Thus, regulation of cytosolic and organelle ATP levels, and ATP/ADP transport across the mitochondrial and plastid envelope, give rise to a particularly complex ATP dynamics in plant cells, that is critically dependent on the tissue type and external conditions (Neuhaus et al., 1997; Flügge, 1998; Reiser et al., 2004; Haferkamp et al., 2011). The exact nature of the interplay between mitochondria and chloroplasts in maintaining cytosolic and nuclear ATP homeostasis, especially under changeable conditions, such as light-dark cycles or varying O_2_/CO_2_ status, has been investigated for decades, predominantly using biochemical techniques or *in vivo* NMR (Bailleul et al., 2015; Gardeström and Igamberdiev, 2016). For example, insight into subcellular adenine nucleotide pools have been possible through rapid membrane filter-based fractionation of leaf protoplasts, revealing a complex and dynamic interplay between the three cell compartments (Lilley et al., 1982; Stitt et al., 1982; Gardeström and Wigge, 1988; Krömer and Heldt, 1991; Krömer et al., 1993). The current consensus is that cytosolic ATP is mainly provided by the mitochondria, both in the dark, but also in the light, when photorespiration can be the main driver of ATP synthesis (Igamberdiev et al., 2001). Little is known, however, about differences between organs, tissues and cells and about the characteristics of their specific responses over time.

Similar challenges apply to other physiological and metabolic parameters, such as pH, free Ca^2+^, potentials of thiol redox couples, and concentrations of small molecules including plant growth regulators. However, development of *in situ* reporters has provided increasingly sophisticated understanding of their *in vivo* behaviour (De Michele et al., 2014; Uslu and Grossmann, 2016). For example, detailed insights into subcellular pH gradients and their dynamics have turned out to play a critical role in membrane transport, protein degradation, and energy and ion homeostasis (Schwarzländer et al., 2012; Luo et al., 2015). Likewise, the spatiotemporal characteristics of free Ca^2+^ transients are central to signalling in stress responses and plant-microbe interactions (Choi et al., 2014b; Keinath et al., 2015). The ability to separately monitor redox potentials of the subcellular glutathione pools has revealed a far more reducing cytosolic redox landscape than previously anticipated, and has led to novel concepts of redox regulation and signalling (Marty et al., 2009; Morgan et al., 2013; Schwarzländer et al., 2016).

Recently, different fluorescent sensor proteins for ATP have been engineered (Berg et al., 2009; Imamura et al., 2009; Kotera et al., 2010; Nakano et al., 2011; Tantama et al., 2013; Yoshida et al., 2016). ‘Perceval’ is based on a single circularly permuted mVenus protein fused to the bacterial regulatory protein GlnK1 from *Methanococcus jannaschii*. Competitive binding of ATP and ADP to GlnK1 result in inverse changes of two excitation maxima to provide a ratiometric readout of ATP:ADP (Berg et al., 2009). The ‘PercevalHR’ variant was obtained by mutagenesis, and has an improved dynamic range of about 4 (Tantama et al., 2013). However, both variants are strongly pH-sensitive, requiring pH measurement and correction for meaningful *in vivo* measurements. By contrast, the ratiometric ATeam sensor family were introduced as far less pH sensitive. ATeam sensors share their overall design with the widely used Förster Resonance Energy Transfer (FRET) sensors of the Cameleon family, making use of the ε-subunit fragment of ATP synthase from *Bacillus* sp. PS3 for reversible ATP binding (Imamura et al., 2009; Kotera et al., 2010). ATP binding to the εsubunit induces a conformational change in the sensor structure modifying the relative orientation of the N- and C-terminal donor and acceptor fluorophores (monomeric super-enhanced cyan fluorescent protein (mseCFP); circularly permuted monomeric Venus (cp173-mVenus), a variant of yellow fluorescent protein), increasing FRET efficiency. Both sensor classes have provided insights into subcellular ATP dynamics of animals (Ando et al., 2012; Tarasov et al., 2012; Li et al., 2015; Merrins et al., 2016). However, to date there is only one report on their use in plants (Hatsugai et al., 2012), and a reliable establishment of fluorescence-based ATP monitoring in plants has been lacking. This is despite the prominent role of ATP in the physiological network of plants, including two ATP-producing organelles and frequently fluctuating environmental conditions that determine the development of their flexible body plan.

In this work we set out to establish ATP sensing in plants. First, we generate Arabidopsis lines expressing the biosensor ATeam1 .03-nD/nA in the cytosol, mitochondrial matrix or the plastid stroma, and demonstrate that plants harbouring the probe in the cytosol or plastids are stable and show no phenotypic change. By contrast, lines expressing mitochondrial sensors are dwarfed, but still viable. Second, we validate the biochemical characteristics of the ATeam 1.03-nD/nA sensor *in vitro* under conditions typically encountered in plant systems. Third, we develop an *ex situ* assay for isolated mitochondria to probe ATP transport and synthesis. Fourth, we map tissue differences and gradients of cytosolic MgATP^2−^ concentrations in living seedlings, including cell-to-cell variation in root hairs that inversely correlates with the rate of growth. Finally, we demonstrate how progressive hypoxia leads to characteristic time-dependent changes in MgATP^2−^ dynamics.

## Results

### Generation of Arabidopsis lines for ATP sensing in the cytosol, chloroplasts and mitochondria

To establish ATP measurements in living plants, we generated Arabidopsis lines expressing ATeam1.03-nD/nA in the cytosol, the chloroplast stroma, and the mitochondrial matrix. We selected at least three independent lines for each compartment based on fluorophore expression, two of which were propagated to homozygosity. Despite expression being driven by a CaMV 35*S* promoter, we did not observe strong sensor silencing in subsequent generations contrary to frequent observations for other sensors (Pei et al., 2000; Deuschle et al., 2006; Chaudhuri et al., 2008; Yang et al., 2010; Jones et al., 2014; Behera et al., 2015; Loro et al., 2016; Schwarzländer et al., 2016). Fluorescence in the peripheral cytoplasm, as well as in trans-vacuolar strands and the nucleoplasm, demonstrated cytosolic expression, whilst co-localisation with the chlorophyll auto-fluorescence confirmed chloroplast expression (Figure 1A). Furthermore, all independent lines for cytosol or plastid expression showed a wild-type-like phenotype at the whole plant level (Figure 1B), which was validated by detailed phenotyping quantifying root length, rosette size, inflorescence height and number of siliques (Figure 1 – figure supplement 2 and 3). By contrast, transformants for mitochondrial expression showed a consistently weaker fluorescence and a strong developmental phenotype (Figure 1A,B; Figure 1 – figure supplement 2). The sensor fluorescence co-localized with the mitochondrial matrix marker MitoTracker, but the organisation of the labelled cell structures was perturbed, suggesting abnormal mitochondria. Nevertheless, transformants flowered after 14 weeks and set seed, allowing their propagation. Despite these observations, we recorded the fluorescence of Venus and CFP in 5-day-old seedlings (Figure 1 – figure supplement 1) and found markedly lower Venus/CFP ratios in mitochondria while cytosol and chloroplasts were similar (Figure 1C; Figure 1 – figure supplement 1).

**Figure 1:**
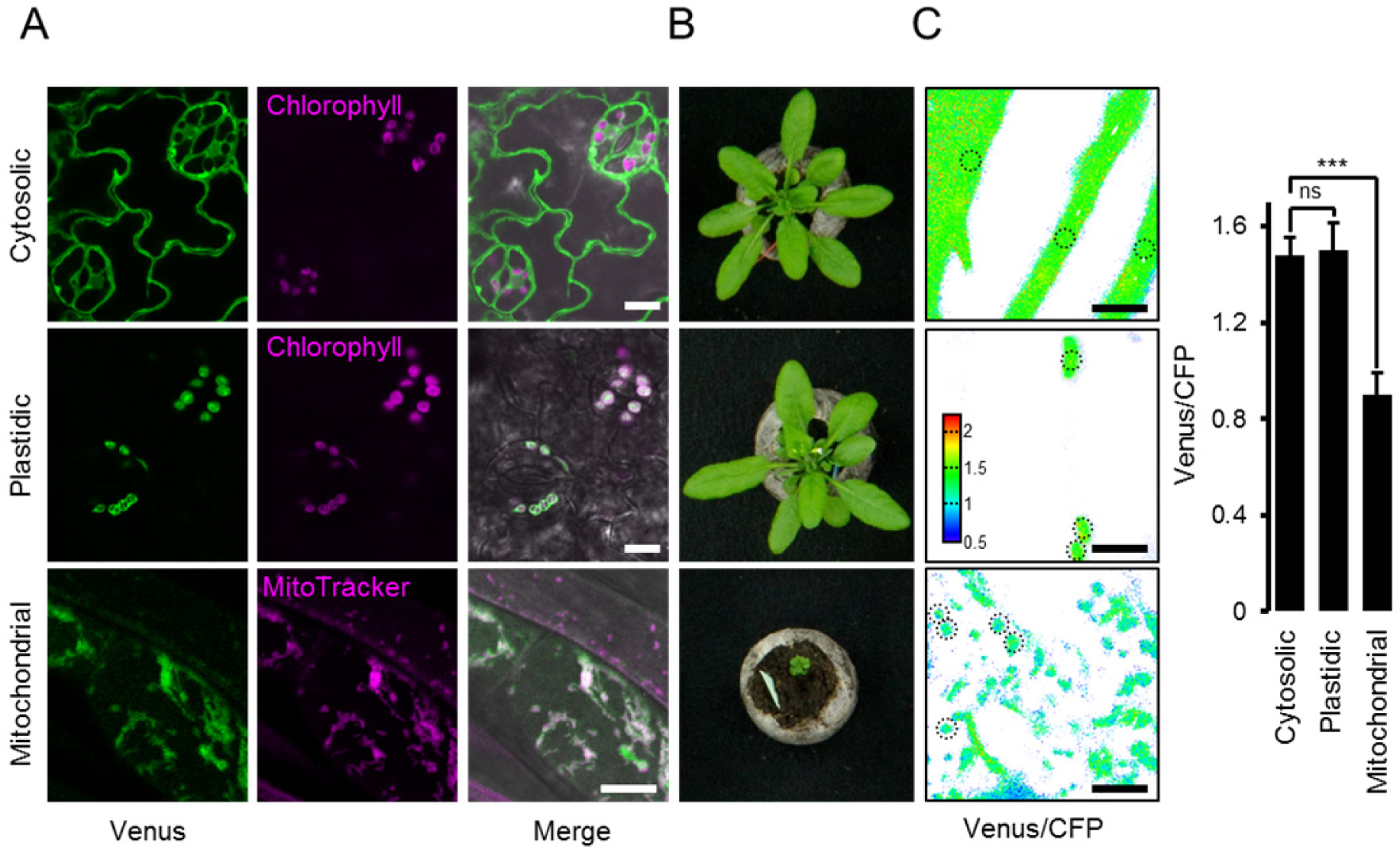
ATeam expression in stable Arabidopsis lines. ATeam1.03-nD/nA was expressed under the control of a 35*S* promoter as unfused protein for localization in the cytosol, as a fusion with the transketolase target peptide (TkTp) for plastid targeting or fused to the *Nicotiana plumbaginifolia* β-ATPase for mitochondrial targeting. (A) Five-day-old seedlings grown vertically on half-strength MS + 1% (w/v) sucrose medium plates were used for CLSM. Venus (green) fluorescence was recorded alongside the chlorophyll fluorescence in cotelydon cells or the mitochondrial marker MitoTracker Orange in cells of the hypocotyl. The merge image shows both fluorescence channels projected on the respective bright field image. (B) Phenotypes of multiple independent lines per construct were compared and a representative image is shown after growth for five weeks on soil. (C) For a ratiometric analysis, fluorescence of Venus and CFP was assessed in hypocotyl cells of 5-day-old seedlings grown as in (A) with power of the 458 nm laser set to 10% (cytosolic and plastidic) and 30% (mitochondrial) of maximal power. Regions of interest (ROIs) of similar size, indicated by dotted lines, were defined to calculate the Venus/CFP ratio shown in the graph. *n* = 36 (cytosol/plastid) or 105 (mitochondria) ROIs in 12 (cytosol/plastid) or 22 (mitochondria) images from 4 (cytosol/plastid) or 6 (mitochondria) individual plants; error bars = SD. ns: *p* > 0.05, *** *p* ≤ 0.001 (*t* test). Scale bar (all panels) = 10 μm.

### In vitro characterisation of purified sensor protein revealing specificity for MgATP^2−^

To reliably interpret *in vivo* measurements in the Arabidopsis sensor lines, we aimed for an in-depth understanding of key sensor characteristics. The K_d_ (ATP), nucleotide specificity and pH sensitivity of the original ATeam family variants were characterised for use in animal cells at 37°C (Imamura et al., 2009). While the newer ATeam1.03-nD/nA variant shows an improved dynamic range (Kotera et al., 2010), its other properties have not been characterised at the required detail, particularly at the pH and temperature conditions likely to be experienced in the plant cytosol, mitochondrial matrix and plastid stroma. We therefore characterised purified ATeam1.03-nD/nA protein (Figure 2A) *in vitro*. The sensor emission spectrum showed a well-defined ratiometric shift in response to ATP. The mseCFP peak (475 nm) decreased and the cp173-mVenus peak (527 nm) increased, with increasing ATP concentrations in the high micromolar/low millimolar range Figure 2B). The isosbestic point was at 512 nm. Purified protein was stable at −86°C and retained the same dynamic range of the freshly purified sensor. However, non-frozen storage caused a decline of sensor responsiveness over time, arising from sensor degradation, probably by proteolysis, explaining the diminished dynamic range by separation of the two chromophores (Figure 2 – figure supplement 1A). Hence, aliquots of purified protein were frozen immediately after purification and stored at −86°C. The ATP response of the sensor, determined at 25°C, was sigmoidal with a spectroscopic dynamic range of 4.0 higher than previously reported at 3.2 (Kotera et al., 2010). The Kd(ATP) was 0.74 mM with a Hill coefficient of 1.02, compatible with a single ATP binding site in the ATP synthase ε-subunit (Yagi et al., 2007) (Figure 2C). The sensor showed no response to ADP and AMP, in agreement with previous reports using other nucleotides, including GTP (Imamura et al., 2009), and confirms the selectivity for ATP. Mg^2+^ titration under saturating ATP showed a strong Mg^2+^ dependence, indicating that the sensor responds selectively to MgATP^2−^, and not to ATP^4−^ (Figure 2D,E). In the absence of ATP, the sensor signals were pH-stable from pH 6.5 to 8.5 (Figure 2F), consistent with the absence of direct pH effects on the fluorophores. Insensitivity to pH was also observed at partially saturating and saturating ATP from pH 7.5 to 8.5, although the spectroscopic sensor response range was diminished below pH 7.0. This decrease in ratio was due to a *bona fide* FRET response (mseCFP donor signal increasing, cp173-mVenus acceptor signal decreasing; Figure 2 – figure supplement 1B), and correlated with the decrease in the MgATP^2−^ species as the pH was lowered. We infer that the sensor response reports the MgATP^2−^ concentration, which itself depends on ambient Mg^2+^ concentration and pH (O’Sullivan and Perrin, 1961; Storer and Cornish-Bowden, 1976; Adolfsen and Moudrianakis, 1978). Since MgATP^2−^ is the species that acts as the cofactor for the large majority of ATP-dependent proteins, the sensor provides a readout of the physiologically significant proportion of the ATP pool that is visible for those proteins. The *MgATP^2−^ response* of the sensor will be referred to as *ATP response* for simplicity in sections of this work.

**Figure 2:**
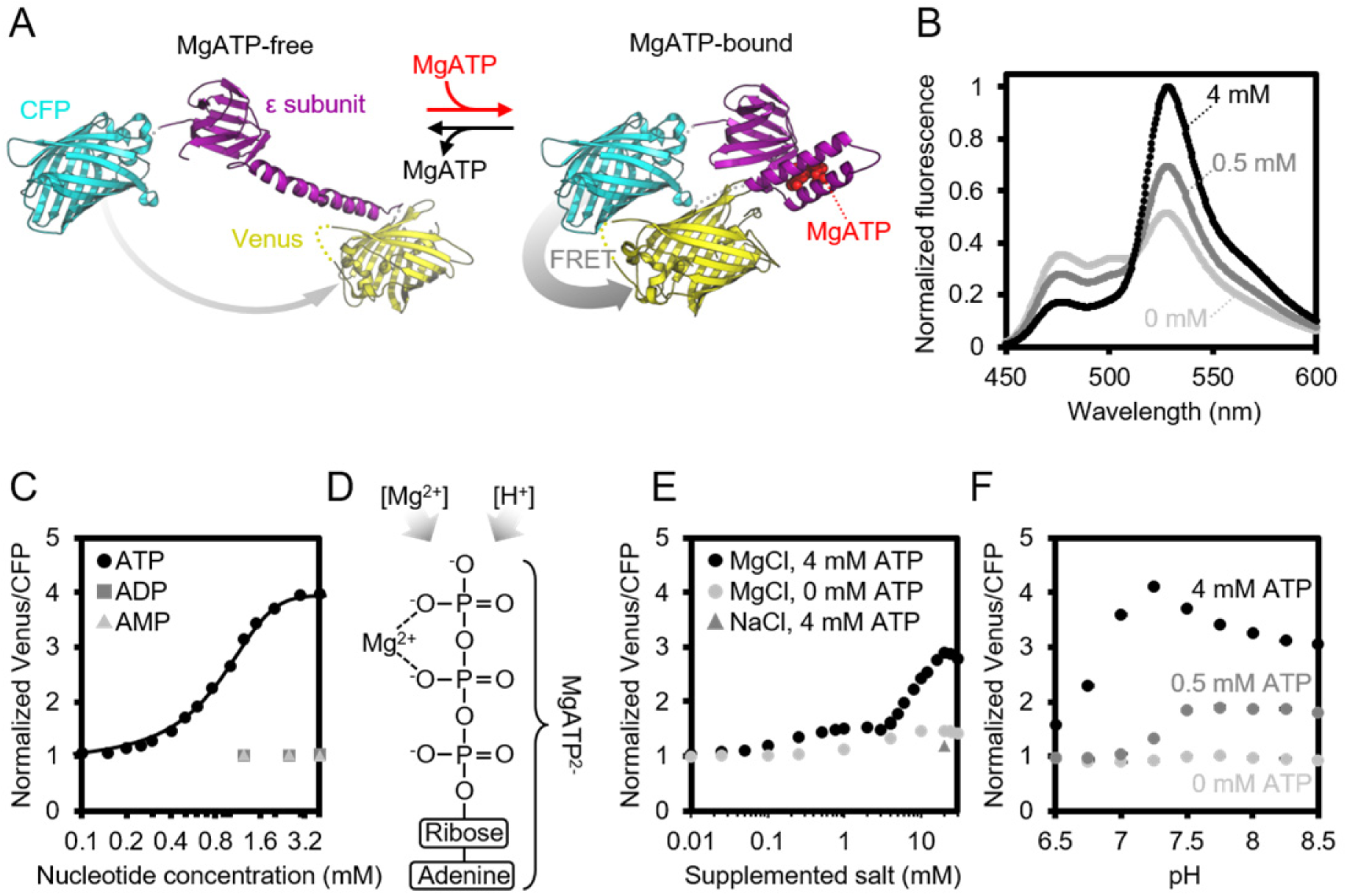
Characteristics of purified ATeam1.03-nD/nA. (A) A cyan fluorescent protein (CFP, PDB: 2WSN) and a variant of the yellow fluorescent protein (Venus, PDB: 3EKJ) were manually linked by the ε-subunit of *Bacillus subtilis* ATP synthase in the MgATP^2−^-bound (PDB: 2E5Y) and MgATP^2−^-free (PDB: 4XD7) state to generate a hypothetical structural model of ATeam. FRET efficiency in the absence or presence of MgATP^2−^ is indicated by a grey arrow. (B) Normalized ATeam emission spectra (excitation at 435 ± 5 nm) in the presence of increasing ATP concentrations and an excess in Mg^2+^ by 2 mM. (C) ATeam was excited at 435 ± 5 nm and the ratio of emission at 527 nm (cp173-Venus) and 475 nm (mseCFP) at 25°C in the presence of adenine nucleotides is plotted. The Boltzmann function was used to fit MgATP^2−^-binding data. (D) Structure of MgATP^2−^. Its stability depends on pH and on the free Mg^2+^ concentration. (E) ATeam Venus/CFP ratios in 4 mM ATP (black points) and 0 mM ATP (grey points) titrated with increasing concentrations of MgCl2. The grey triangle shows the Venus/CFP ratio in the presence of 4 mM ATP and 20 mM NaCl. (F) ATeam Venus/CFP ratios at different pH and in the presence of 0 (light grey), 0.5 (dark grey) and 4 mM (black) MgATP. Data in (B), (C) and (F) is averaged from four technical replicates and error bars are represented as SD, but too small to be displayed.

### Establishing an ex situ assay to monitor mitochondrial ATP dynamics

*In vivo* ATP dynamics in the cell are governed by production and consumption. On the production side, mitochondria export ATP to the cytosol (and the nucleus in turn by diffusion), facilitated by oxidative phosphorylation at the matrix surface of the inner mitochondrial membrane and membrane gradient-driven ATP extrusion by a very active ADP/ATP carrier (AAC) system (Haferkamp et al., 2011; Gout et al., 2014). We hypothesized that supplementing purified, functional mitochondria with external sensor protein would allow monitoring ATP transport fluxes into and out of the mitochondria, and also dissect the role of adenylate kinase (AK), which is thought to be localised to the intermembrane space (Figure 3A). Such a system would remove the influence of other cellular processes that impact on MgATP^2−^ concentration in the cytosol, including ATP hydrolysis, transport across membranes of other cell compartments, as well as changes in pH and Mg^2+^ concentrations.

**Figure 3:**
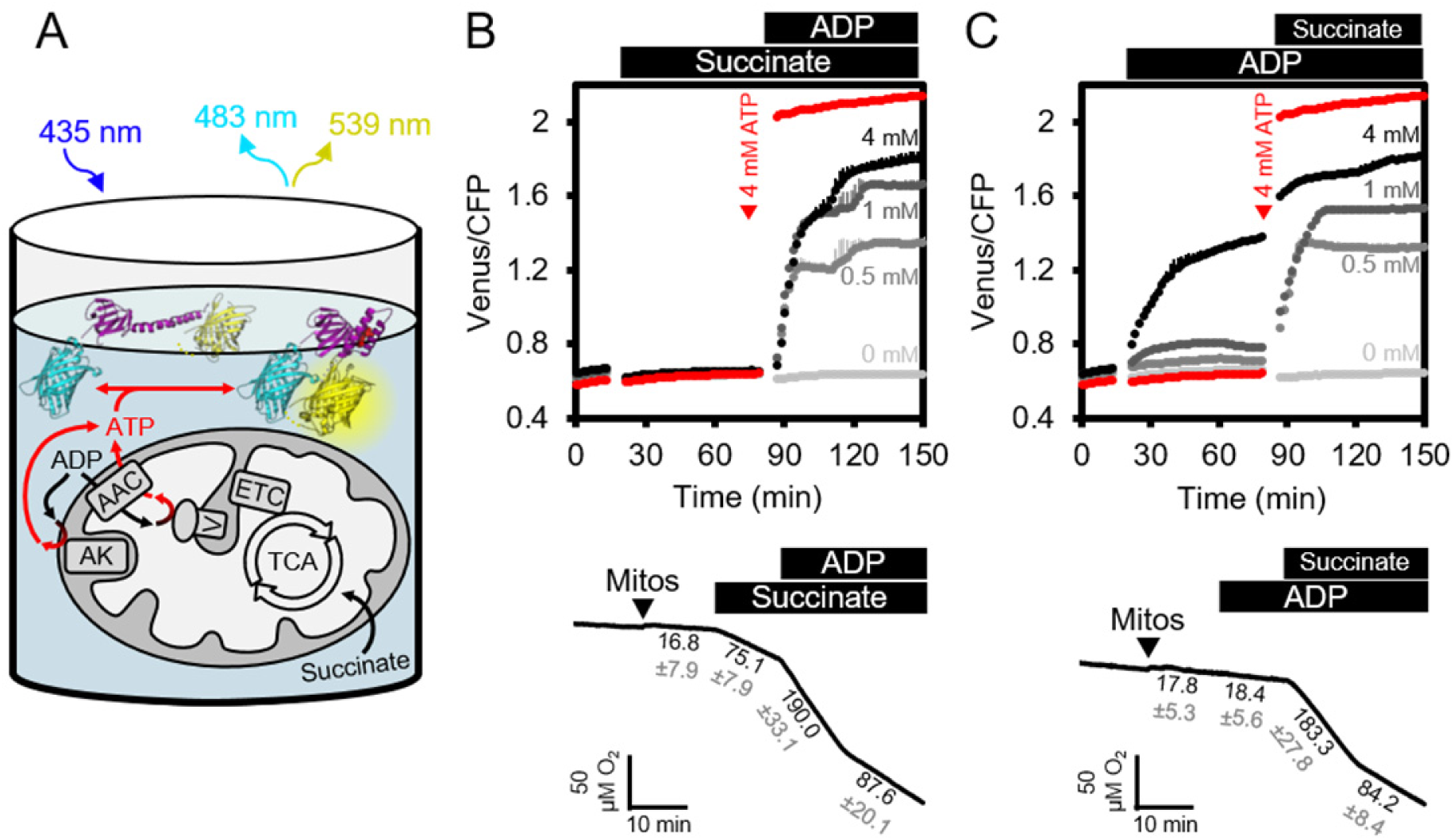
ATP fluxes in isolated Arabidopsis mitochondria. (A) Pure intact mitochondria isolated from 2-week-old Arabidopsis seedlings were mixed with basic incubation medium and purified ATeam in 96-well microtiter plates. Mitochondria were fed with succinate as a respiratory substrate to fuel the tricarboxylic acid (TCA) cycle and the electron transport chain (ETC). ATP, chemiosmotically generated by ATP synthase (complex V) in the matrix, is exchanged for ADP through the ADP/ATP carrier (AAC), while ATP generated by adenylate kinase (AK) in the intermembrane space does not require transport across the inner mitochondrial membrane. Extramitochondrial ATP exported from the mitochondria is sensed by ATeam in the medium as MgATP^2−^. (B, C) ATeam was used as shown in (A) and 10 mM succinate was added either before (B) or after (C) ADP at concentrations between 0 and 4 mM. ATP was added to 4 mM as reference at the indicated time point (red traces). *n* (technical replicates) = 4; error bars = SD. Lower panels show polarographic oxygen consumption assays performed with a Clark-type electrode in parallel. Mitochondria, succinate and ADP were added to the basic incubation medium as indicated. A representative trace from an individual experiment is shown and oxygen consumption rates (nmol min^-1^ mg^-1^ protein) for each respiratory state are given as mean ± SD from three technical replicates.

We supplemented freshly isolated Arabidopsis seedling mitochondria with ATeam sensor protein using the same medium as used for *in vitro* sensor characterisation (Figure 3B). The medium contained Mg^2+^ and Pi in excess, and was set to pH 7.5 to resemble cytosolic pH (Ratcliffe, 1997; Schulte et al., 2006). The exact concentration of the sensor is not critical, because the ratiometric FRET readout is self-normalizing, although very low or very high sensor concentrations were avoided, to prevent low signal-to-noise, or ATP-buffering by the sensor itself when close to its K_d_(ATP).

The FRET ratio did not change on addition of succinate as respiratory substrate in the absence of ADP, but subsequent addition of ADP led to an increase in FRET that plateaued at a steady state value depending on the ADP concentration (Figure 3B). Polarographic oxygen consumption assays performed in parallel confirmed respiratory activity responses and coupling of the mitochondria with a respiratory control coefficient of around 2 (Figure 3B), and confirmed that the sensor responded rapidly to ATP generated by respiring mitochondria and exchanged by the AAC. Nevertheless, addition of ADP before succinate revealed that ADP alone was sufficient to cause the sensor to respond in a dose-dependent manner, although with smaller FRET increases (Figure 3C). Considering the sensor does not respond to ADP (see Figure 2C), the response is indicative of ATP production in the absence of active respiration. FRET ratios increased further after subsequent addition of succinate, reaching a similar plateau value to before. Production of ATP from ADP alone in the absence of a respiratory substrate ruled out ATP synthase activity, but would be consistent with AK activity leading to conversion of ADP to ATP and AMP (Busch and Ninnemann, 1996). This implies that the assay provides an integrated readout of the combined activities of ATP synthase, AK and the AAC.

### Dissection of mitochondrial ATP production, ATP/ADP exchange, and adenylate kinase activity

The mitochondrial assay allows investigation of the role of AK and AAC in controlling mitochondrial ATP dynamics. To test whether AK is involved in ATP production in the absence of respiratory electron transport, we predicted that ATP production would be sensitive to the presence of AMP driven by mass action, in a fully reversible reaction catalysed by AK (Figure 4A). Consistent with this view, addition of AMP following the increase in FRET ratio triggered by ADP, led to a gradual, dose-dependent decrease of FRET (Figure 4B). Likewise, the presence of AMP before addition of ADP, inhibited ADP-induced ATP generation (Figure 4C).

**Figure 4:**
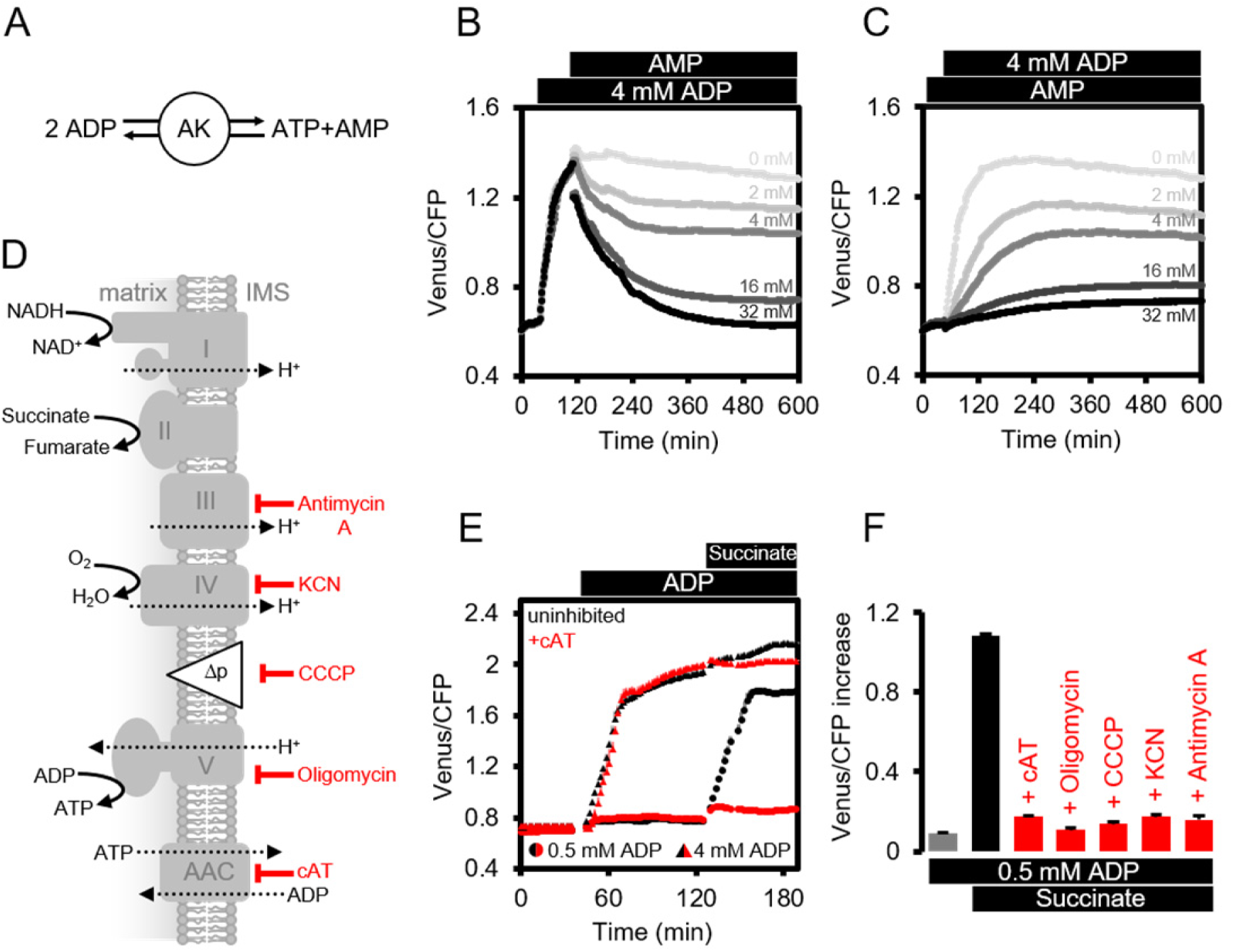
Modulating the ATP production in isolated Arabidopsis mitochondria. (A) Reaction catalyzed by adenylate kinase (AK). (B, C) ATeam was used in a setup as shown in Fig. 3A. Isolated mitochondria and purified ATeam were mixed with AMP between 0 (light grey) and 32 (black) mM either after (B) or before (C) the addition of 4 mM ADP and the ATeam Venus/CFP ratio was recorded. *n* (technical replicates) = 4. (D) Representation of the mitochondrial electron transport chain (complexes I-IV) that generates a proton motive force (Δp) used by the ATP synthase (complex V) to produce ATP. The ADP/ATP carrier (AAC) transports ATP from the mitochondrial matrix to the intermembrane space (IMS) in exchange for ADP. Treatments that diminish mitochondrial ATP production or transport and their site of action are indicated. (E) Untreated mitochondria (black symbols) or mitochondria treated with 10 μM cAT (red symbols) were fed with ADP at 0.5 mM (circles) or 4 mM (triangles) followed by 10 mM succinate. (F) The inhibitory effect of treatments summarized in (F) on mitochondrial ATP production was calculated through the Venus/CFP increase after addition of ADP (grey), ADP and succinate (black) or ADP and succinate under inhibition (red). cAT, carboxyatractyloside; CCCP, carbonyl cyanide m-chlorophenyl hydrazone; KCN, potassium cyanide. E and F show the mean of three technical replicates; error bars = SD.

To test the role of AAC-mediated ADP/ATP exchange across the inner membrane (Figure 4D), we added carboxyatractyloside (cAT) to block the AAC (Figure 4E). Addition of 4 mM ADP in the absence of respiratory activity resulted in a similar FRET increase in both control and cAT-treated mitochondria, suggesting that AK-derived ATP did not rely on AAC-mediated ADP or ATP transport. Subsequent energization by succinate lead to a slight but reproducible further increase in the control, which was absent for the cAT-treated mitochondria. This difference shows that matrix-exposed ATP synthase cannot contribute in the presence of cAT and validates effective AAC inhibition. At lower concentrations of ADP (0.5 mM), when ATP production by AK is close to the detection limit (see Figure 3C), succinate gave an increase in detectable MgATP^2−^ in the absence of cAT, but this was almost completely abolished in the presence of cAT (Figure 4E). This is consistent with localisation of AK outside the inner mitochondrial membrane and supports localisation in the mitochondrial intermembrane space, as reported previously for other plant species (Day et al., 1979; Birkenhead et al., 1982; Stitt et al., 1982; Roberts et al., 1997; Zancani et al., 2001), but contrasts the situation in mammals where AK3 isoforms are also present in the matrix (Schulz, 1987).

By exploiting the differential sensitivity of AK and the AAC/ATP synthase system to ADP, it is possible to minimise the contribution of AK to ATP production, and use low ADP concentrations (0.5 mM) to selectively monitor ATP produced by ATP synthases and AAC activity (Figure 4C). ATP synthesis then strictly depends on ADP import, ATP synthase functionality, the proton motive force and functional electron transport (Figure 4F). Treatment with specific inhibitors that cover those four functional levels consistently prevented the FRET increase in response to mitochondrial energization by succinate (Figure 4D-F), validating the assay as a means to monitor functional changes at specific steps in bioenergetic pathways.

### High-throughput measurements of MgATP^2−^ in planta by fluorimetry

To allow for high-throughput measurements of MgATP^2−^ levels *in planta*, we optimized the microtiter plate fluorimetry setup for plant tissues and whole seedlings. Ratiometric analysis makes measurements independent of sensor expression level, as well as tissue amount and shape, provided there is sufficient signal-to-noise and little interference from tissue auto-fluorescence. Emission spectra were recorded with excitation at 435 nm from 7-day-old intact Arabidopsis seedlings (Sweetlove et al., 2007), and leaf disks of 4-week-old plants expressing cytosolic ATeam (Figure 5A,B). Both sample types showed fluorescence spectra that were practically identical with the purified sensor protein, while auto-fluorescence was low by comparison. Chlorophyll fluorescence was effectively separated and did not cause any significant interference. This was independently confirmed also for the chloroplastic sensor by confocal microscopy at the individual chloroplast level (Figure 5 – figure supplement 2).

**Figure 5:**
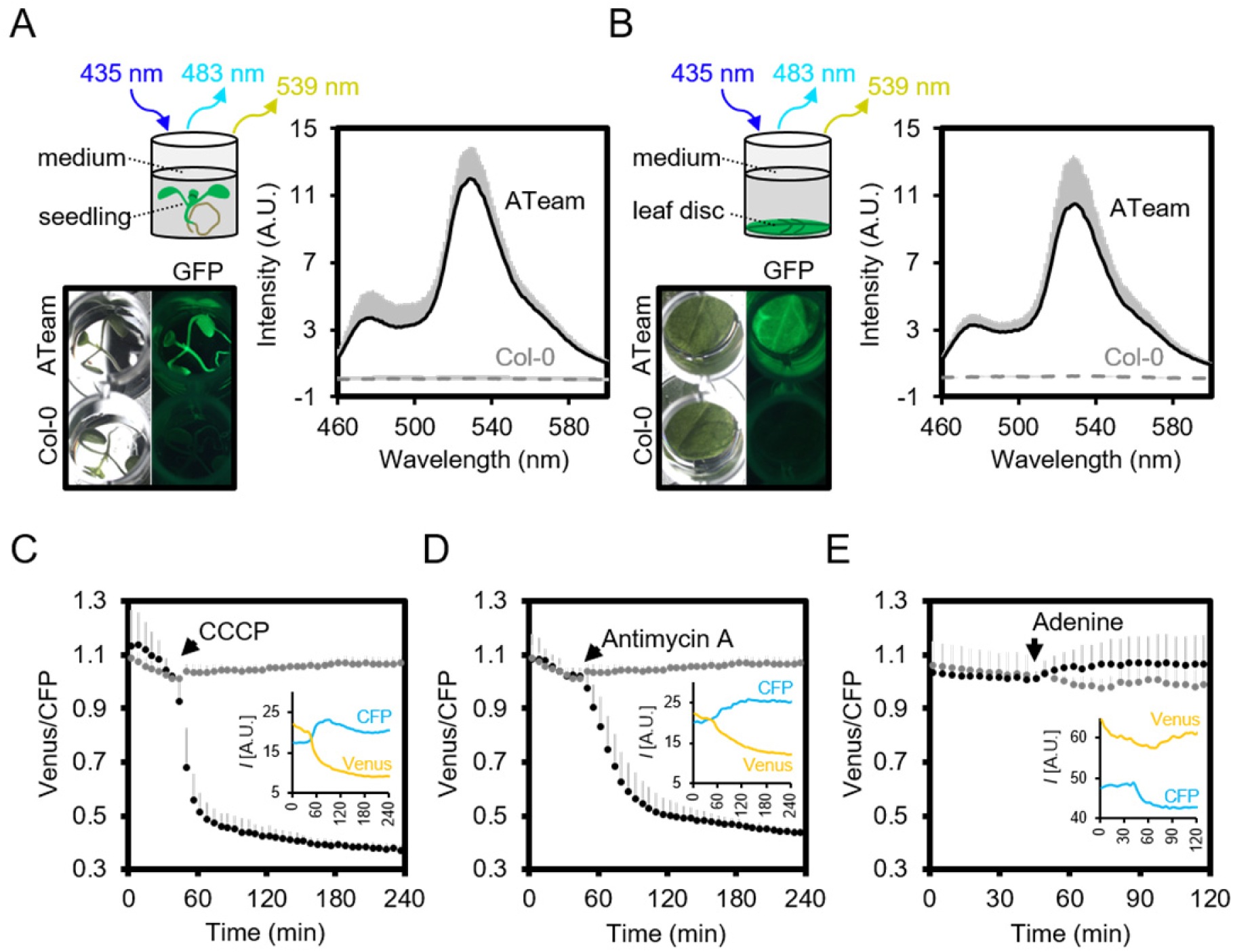
Fluorimetry setup to monitor MgATP^2−^ dynamics *in planta*. (A,B) Seven-day-old Arabidopsis seedlings (two per well; A) or leaf discs of individual 4-w-old plants (B) were submerged in imaging medium on 96-well microtiter plates. Fluorescence of plant material stably expressing cytosolic ATeam was checked with an epifluorescence microscope equipped with a GFP filter and compared to wild-type Col-0. Fluorescence emission spectra between 460 and 600 nm were recorded using a plate reader and an excitation wavelength of 435 ± 10 nm. *n* = 5; error bars = SD. (C-E) Per well, two 7-day-old Arabidopsis seedlings expressing no sensor (Col-0) or cytosolic ATeam were excited at 435 ± 10 nm and the emission at 483 ± 9 nm (mseCFP) and 539 ± 6.5 nm (cp173-Venus) was recorded. CCCP (100 μM), antimycin A (100 μM) or adenine (10 mM) were added where indicated (black data points) while control plants were left untreated (grey data points). Emission in wells with Col-0 plants was averaged and subtracted from that of ATeamexpressing plants to correct for background fluorescence. Data shown and used for background subtraction is the average of 3-4 wells and error bars are SD. Insets show the fluorescence emission intensity (*I*) of Venus and CFP in representative individual wells.

To assess the speed and range of the sensor response *in vivo*, we monitored the FRET ratio of seedlings in microtiter plates over time in the presence and absence of carbonyl cyanide m-chlorophenyl hydrazone (CCCP), antimycin A, and adenine, to modify endogenous MgATP^2−^ concentrations. CCCP dissipates proton gradients over cell membranes, inhibiting ATP production and increasing ATP consumption; antimycin A inhibits mitochondrial electron transport at complex III (Figure 4D); while adenine feeding has been demonstrated to lead to an increase of the cellular MgATP^2−^ concentration by acting as a substrate for ATP synthesis (Loef et al., 2001; Gout et al., 2014). We focused on the cytosol as an integration space for ATP fluxes from and to other subcellular locations, where steady-state concentration of MgATP^2−^ is set by the interplay of numerous synthesis, hydrolysis and transport processes.

CCCP and antimycin A both triggered a rapid and pronounced decrease in FRET, while adenine led to a modest, but reproducible increase (Figure 5C,D,E; Figure 5 – figure supplement 1). Varying the medium pH between 6.0 and 8.5 showed that the decrease in FRET after antimycin A treatment was independent of pH. Also after CCCP addition, only a small fraction of the response was due to cytosolic acidification, and destabilization MgATP^2−^ in turn (Figure 5 – figure supplement 3). The maximal spectroscopic response range was about 3, slightly lower than the range *in vitro*, which is partially accounted for by the excitation wavelength and emission bandwidth used. We infer that the sensor was intact and functional *in vivo*. High, but not fully saturated FRET at steady state allowed an estimate of cytosolic MgATP^2−^ concentrations in the range of about 2 mM, averaged over seedling tissues. This is consistent with previous estimations and textbook values (Taiz et al., 2015), but higher than reported in other cases (Gout et al., 2014).

### A MgATP^2−^ map of the Arabidopsis seedling

Heterogeneity and gradients between tissues and cells have been of major interest in plant hormone signalling and development, and fluorescent sensors for abscisic acid and auxin were introduced recently (Brunoud et al., 2012; Wend et al., 2013; Jones et al., 2014; Waadt et al., 2014). Analogous insights are largely lacking for metabolites and co-factors, despite the fact that metabolism underpins development (Sweetlove et al., 2017), and that abiotic factors, such as hypoxia, act as signals (Considine et al., 2017). To measure potential differences in cytosolic MgATP^2−^ concentrations between tissues and cells *in vivo*, we performed a confocal microscopy analysis of intact 5-day-old Arabidopsis seedlings (Figure 6A). The seedlings were kept in the dark for 30 min before image acquisition to avoid potential effects of active photosynthesis. An overview of FRET across the seedlings revealed large tissue differences. Cotyledons displayed high values, which were lower in the hypocotyl and dropped abruptly at the shoot-root transition. The root showed low values which then increased in the root tip. The relative differences were reproducibly observed between individual seedlings, and consistent in independent sensor lines (#1.1, #3.6; Figure 6B; Figure 6 – figure supplement 1). The area around the shoot apex showed low ratios, comparable with those of the root, in many but not all individuals. By contrast, differences between tissues were less pronounced in etiolated 5-day-old seedlings under identical conditions (Figure 6C,D), with comparatively lower FRET in the shoots and higher FRET in the roots. The steep gradient at the shoot-root transition was also absent, indicating that photo-morphogenesis is required for its formation.

**Figure 6:**
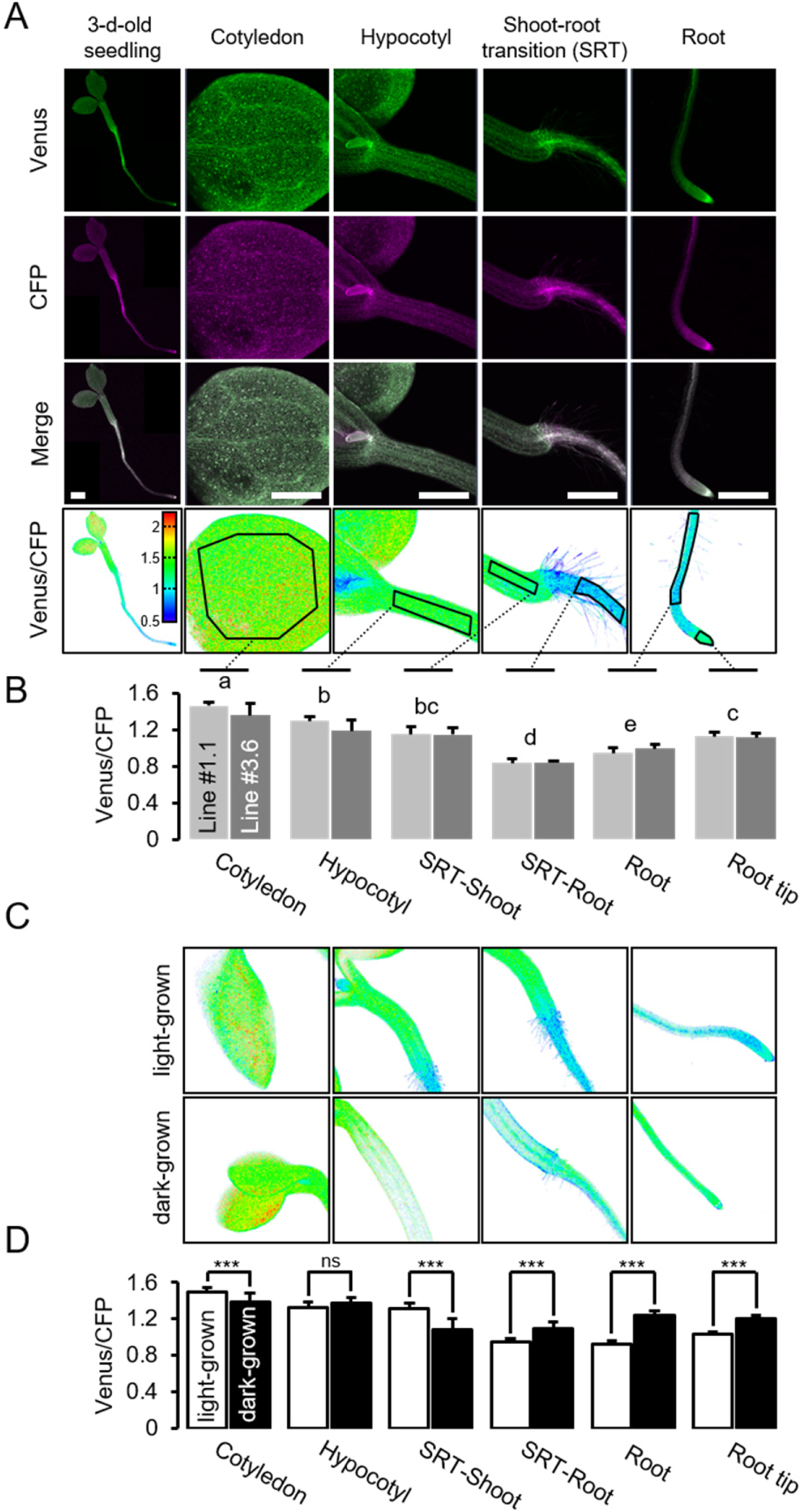
A MgATP^2−^ map of the Arabidopsis seedling. (A) To map the MgATP^2−^ levels in tissues of Arabidopsis seedlings, 3-day-old (whole seedling) or 5-day-old (close-ups) plants expressing cytosolic ATeam were analysed by CLSM. Fluorescence of Venus (green) and CFP (magenta) was recorded and the ratio is plotted as a false-color image where high Venus/CFP values (red) correspond to high MgATP^2−^ levels. In the close-ups, Venus/CFP ratios were analysed in the indicated regions of interest. Scale bar = 500 μm. (B) Graphs represent data from two independent lines. *n* (per line) = 6; error bars = SD. Data from both lines were pooled for a statistical analysis with a one-way ANOVA followed by the Tukey test (*p* ≤ 0.05) and different letters indicate significant differences. The experiment was repeated two times with consistent results. (C, D) CLSM analysis of 5-day-old seedlings either grown in the light or etiolated in the dark. SRT, shoot-root transition. *n* (seedlings per condition) = 11; error bars = SD. ns: *p* > 0.05, *** *p* ≤ 0.001 (two-way ANOVA followed by the Tukey test)).

To test whether the ratios observed could be attributed to MgATP^2−^ concentrations rather than optical artefacts from the different tissue geometries, or other biochemical modifications of the sensors, CCCP was used to deplete MgATP^2−^. In addition, the medium buffer pH was set to 7.5 to avoid destabilisation of MgATP^2−^ as a result of cytosolic acidification. In the presence of CCCP, FRET ratios decreased towards a similar minimum value in all tissues and tissue heterogeneity was gradually abolished (Figure 6 – figure supplement 2A,B). Tissue-specific kinetics that could be resolved by CLSM may reflect differential capacity of tissues to maintain MgATP^2−^ concentrations or simply differential penetration by CCCP. To independently validate the overall difference between shoot and root, we used the microtiter well fluorimetry approach to measure shoot and root samples separately, as attempted previously in tissue extracts (Mustroph et al., 2006). The full spectra indicated *bona fide* FRET differences, which were abolished by CCCP treatment (Figure 6 – figure supplement 2C,D). It is important to note that relative and normalized FRET changes, but not absolute FRET values, can be compared between different fluorimetry/microscopy setups due to different excitation and emission detector configurations.

The conclusion that *in vivo* FRET heterogeneity reliably registered MgATP^2−^ heterogeneity, prompted us to further resolve differences at the cellular level. Comparing pavement and guard cells of abaxial cotyledon epidermis by a region of interest (ROI) analysis did not reveal any differences in our hands (Figure 7A). A previous report from older, true leaves had shown higher MgATP^2−^ concentrations in guard cells than in pavement cells, suggesting that differences can be induced depending on developmental and environmental conditions (Hatsugai et al., 2012). Analysis of seven cell layers of the shoot-root transition zone revealed a continuous gradient, indicating that neighbouring cells can maintain different, but stable, MgATP^2−^ gradients (Figure 7B). At the root tip, cap cells showed low MgATP^2−^. Similarly, low FRET values were observed at wounding sites (Figure 7C). A hotspot of high MgATP^2−^ levels was localized in the columella just below the quiescent centre (Figure 7D).

**Figure 7:**
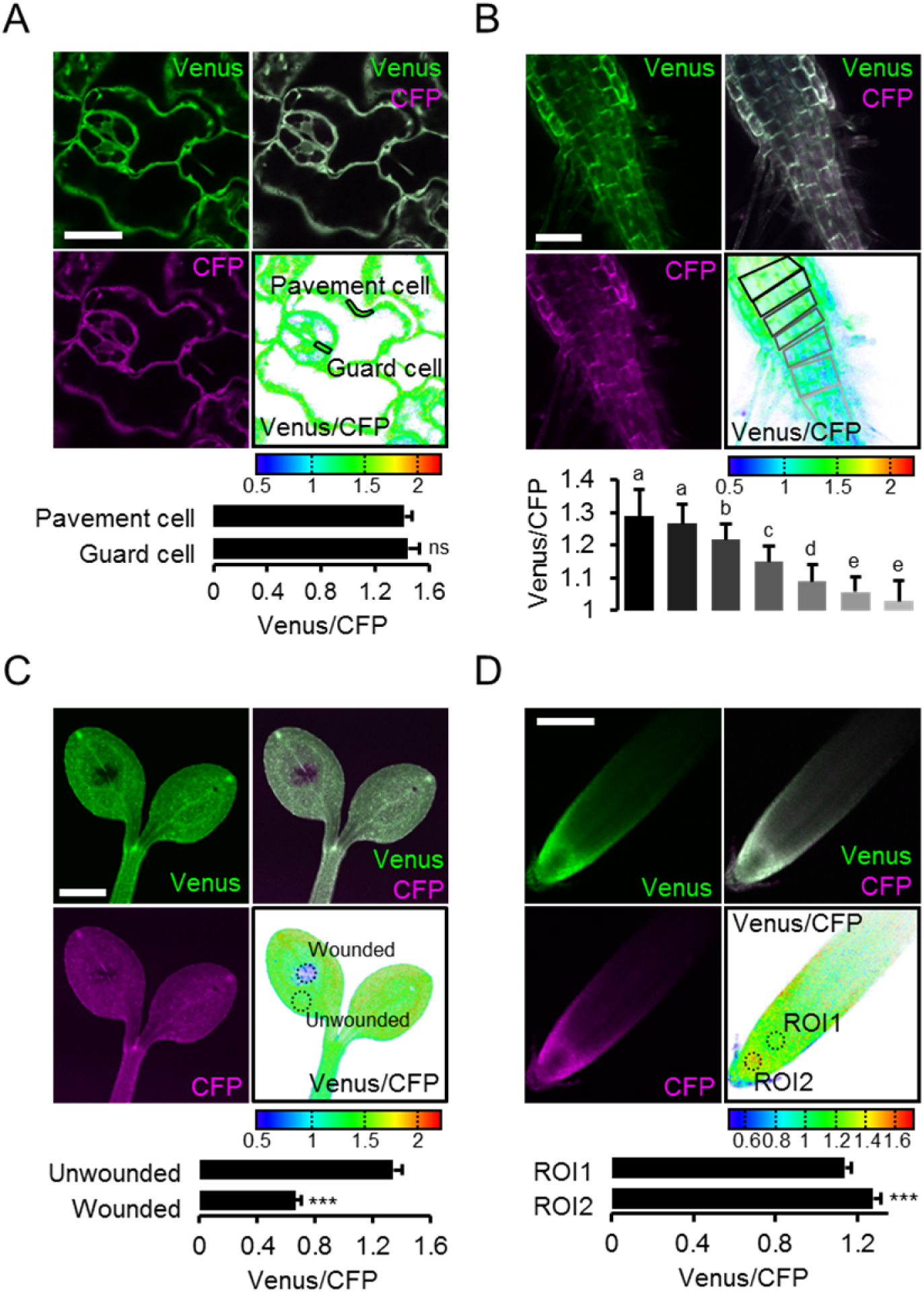
Local MgATP^2−^ heterogeneity in Arabidopsis seedling cells and tissues. Fluorescence of Venus (green) and CFP (magenta) from cytosolic ATeam was recorded by CLSM in 5-day-old Arabidopsis seedlings. The ratio is plotted as a false-color image where high Venus/CFP values (red) correspond to high MgATP^2−^ levels. (A) Venus/CFP ratios were assessed in indicated regions of interest to compare guard cells and pavement cells of the cotyledon epidermis. *n* = 24 pairs of pavement and guard cell from 5 individual plants; error bars = SD. ns: *p* > 0.05 (*t* test). Scale bar = 20 μm. (B) Venus/CFP ratios were assessed in indicated cell layers at the shoot-root transition. Region-of-interest analysis of successive cell layers is indicated in gray scale. *n* = 11; error bars = SD. Different letters indicate statistical differences in a one-way ANOVA followed by the Tukey test (p ≤ 0.05). Scale bar = 100 μm. (C) Cotyledons were wounded with a needle and Venus/CFP ratios were assessed in the indicated regions of interest representing wounded or unwounded tissue. *n* = 5; error bars = SD, *** *p* ≤ 0.001 (*t* test). Scale bar = 500 μm. (D) The Venus/CFP ratio was assessed in two regions of interest at the root tip. *n* = 12; error bars = SD, *** *p* ≤ 0.001 (*t* test). Scale bar = 100 μm.

### Root hair growth speed is correlated with MgATP^2−^ levels as indicated by light sheet fluorescence microscopy

High respiration has been found during pollen tube growth (Dickinson, 1965) and the same is likely for other rapidly tip-growing cells, such as root hairs. On the one hand, high respiration rates may give rise to high ATP levels; on the other, fast growth rates may nevertheless cause ATP depletion. Addressing the question of the relationship between ATP levels and growth of single cells has been technically impossible. We employed light sheet fluorescence microscopy (LSFM) for 4D imaging of the growing Arabidopsis root to quantify growth rates and FRET of the individual root hairs (Figure 8A). We measured both growth speed and FRET ratio of individual hair cells (Figure 8B). While the most rapidly growing 20% of hair cells elongated eight times more quickly than the slowest 20%, the latter showed an increase in their FRET ratio indicating that their cytosolic MgATP^2−^ concentration was increased (Figure 8C,D). The average growth speed was inversely correlated with cytosolic MgATP^2−^ content and lowered MgATP^2−^ was exclusively detected in rapidly growing hair cells (Figure 8E). Interestingly, there was no evidence for oscillations in MgATP^2−^ levels similar to those observed for Ca^2+^ (Hepler et al., 2001; Candeo et al., 2017), as FRET ratios remained stable and did not show any periodic oscillations in the order of seconds, as validated by Fourier analysis (Figure 9).

**Figure 8:**
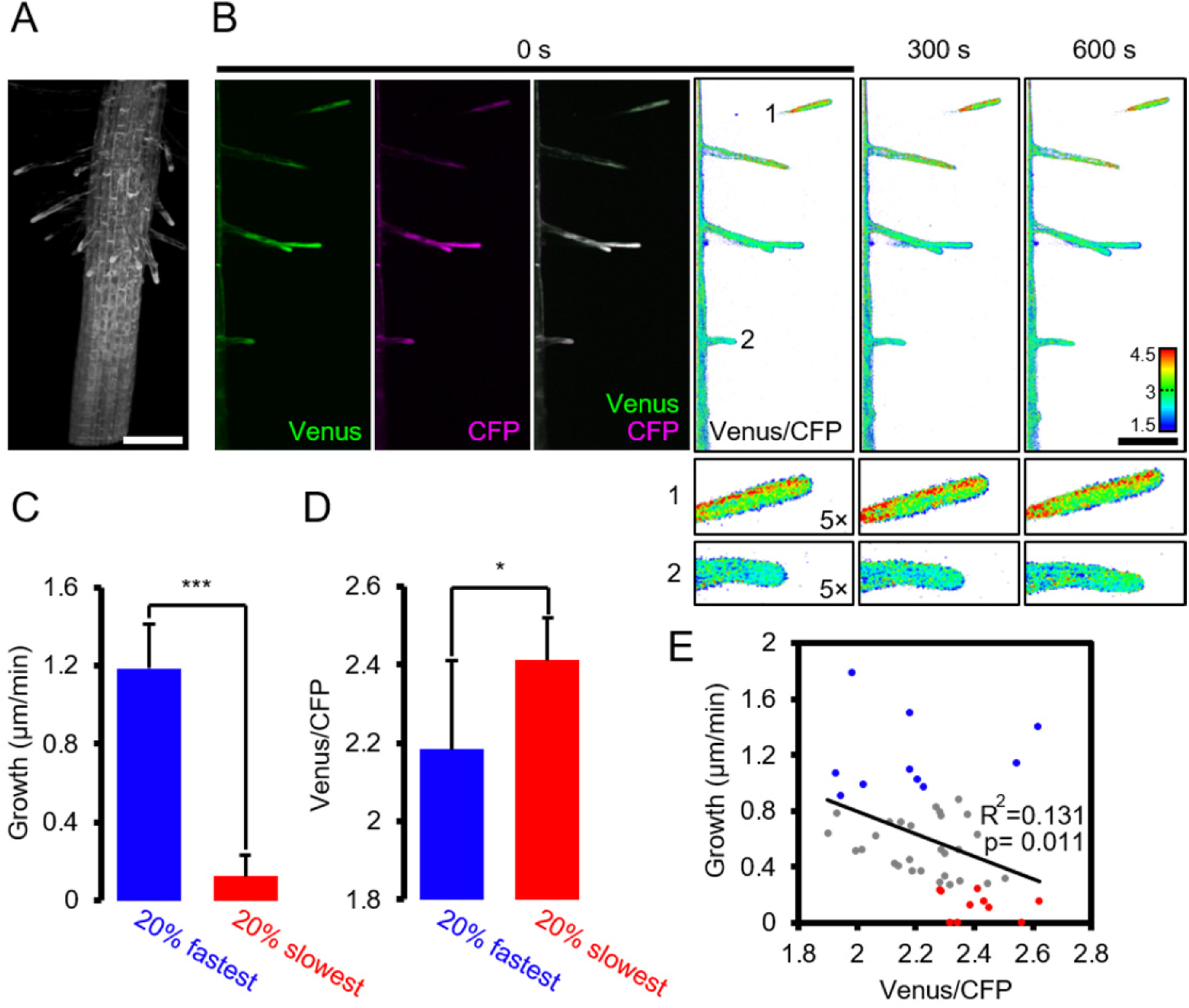
Light sheet fluorescence microscopy (LSFM) analysis of MgATP^2−^ levels in growing root hair cells. Hair cell growth on roots from 6-day-old seedlings expressing cytosolic ATeam was followed for 10 min by LSFM. (A) Maximum projection of the Venus signal in the elongation and lower maturation zone generated from 4D *in vivo* imaging data of the growing Arabidopsis root. Scale bar = 100 μm. (B) Representative root area with hair cells in different developmental stages. Fluorescence of Venus (green) and CFP (magenta) was recorded and the ratio is plotted as false-color images over three time points. Scale bars = 100 μm. Five times magnified image sections exemplify (1) a slow-growing and (2) a fast-growing hair cell. (C) Hair cells in the top and bottom 20% quantile interval of growth speed and (D) the corresponding cytosolic ATeam FRET ratio measured in the tip of the same cells. *n* = 48 hair cells from 6 roots; error bars = SD, * *p* ≤ 0.05, *** *p* ≤ 0.001 (*t* test). (E) Growth speed of individual hair cells as function of their cytosolic ATeam FRET ratio. Values of the slowest and fastest growing cells as included in (C) and (D) are indicated in red and blue; line indicates linear regression. Significance of correlation, expressed as a p-value determined by an *F* test, and the coefficient of determination (R^2^) based on all data points are provided.

**Figure 9:**
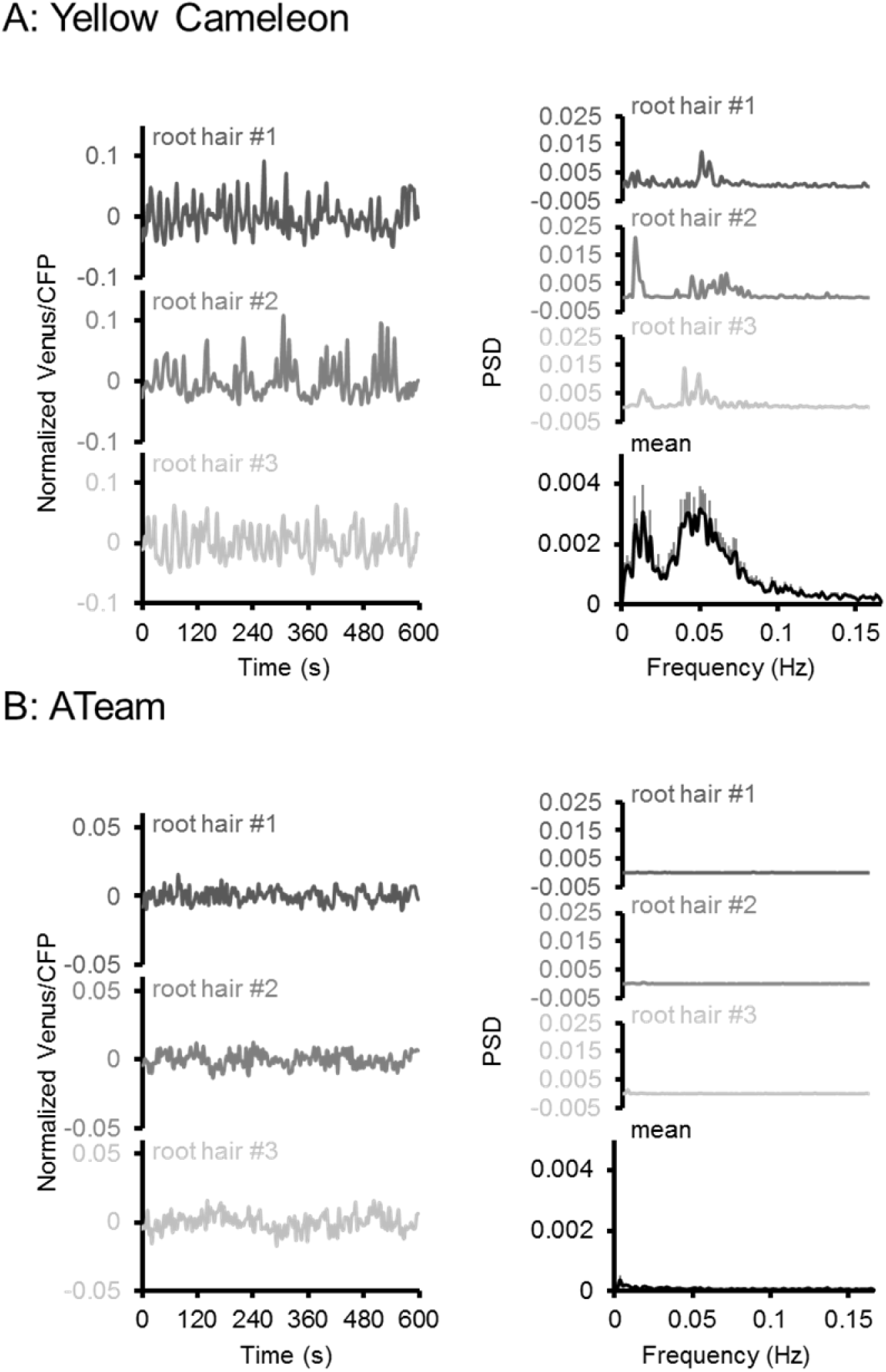
Oscillation analysis in root hair cells. FRET analyses in root hair tips of 6-day-old seedlings expressing (A) NES-Yellow Cameleon 3.6 and (B) cytosolic ATeam1.03-nD/nA by LSFM. Oscillations of three representative root hairs are shown as normalized Venus/CFP ratios and Power Spectral Density (PSD) spectra by Fourier analysis. The mean PSD spectra represent 22 root hairs each. Error bars = SEM. Note the presence of typical Ca^2+^ oscillations around 0.05 Hz in the Cameleon dataset, which were absent in the ATeam dataset.

### Live monitoring of the decline in cytosolic MgATP^2−^ levels during hypoxia

Several stress conditions have been correlated with a cellular energy crisis. Hypoxia has a direct impact on mitochondrial ATP production, since lack of oxygen as final electron acceptor inhibits the respiratory chain. However, flexible metabolic responses have been described in response to hypoxia to prolong maintenance of cellular energy supply (Geigenberger et al., 2000; Geigenberger, 2003; van Dongen et al., 2009; Zabalza et al., 2009; van Dongen and Licausi, 2015). To monitor MgATP^2−^ dynamics during the progression of hypoxia, we used oxygen-proof, transparent tape to seal medium-filled wells containing individual seedlings grown on agar plates and in liquid culture, with no residual air space (Figure 10; Figure 10 – figure supplement 1). The experiments were carried out in the dark (except for excitation flashes for fluorimetric readings) to avoid oxygen evolution by photosynthesis. Seedlings in non-sealed wells served as controls. All FRET ratios showed an initial decrease immediately after immersion. This was followed by a phase of steady decrease for sealed seedlings, while unsealed seedlings retained FRET ratios around the starting values. Re-oxygenation by removal of the seal allowed full recovery to control values, while maintaining the seal led to a phase of sharp decrease and plateauing of FRET ratios at low values, similar to those seen for CCCP treatment. While the response characteristics were reproducible, the onset and rates of the different phases varied between individual seedlings, probably as a result of differences in biomass, respiration rate and the rate of oxygen decrease in turn. The hypoxia model demonstrates the possibility to reliably monitor subcellular MgATP^2−^ dynamics live during stress insult, and may be flexibly adjusted to other external conditions, tissues and genetic backgrounds.

**Figure 10:**
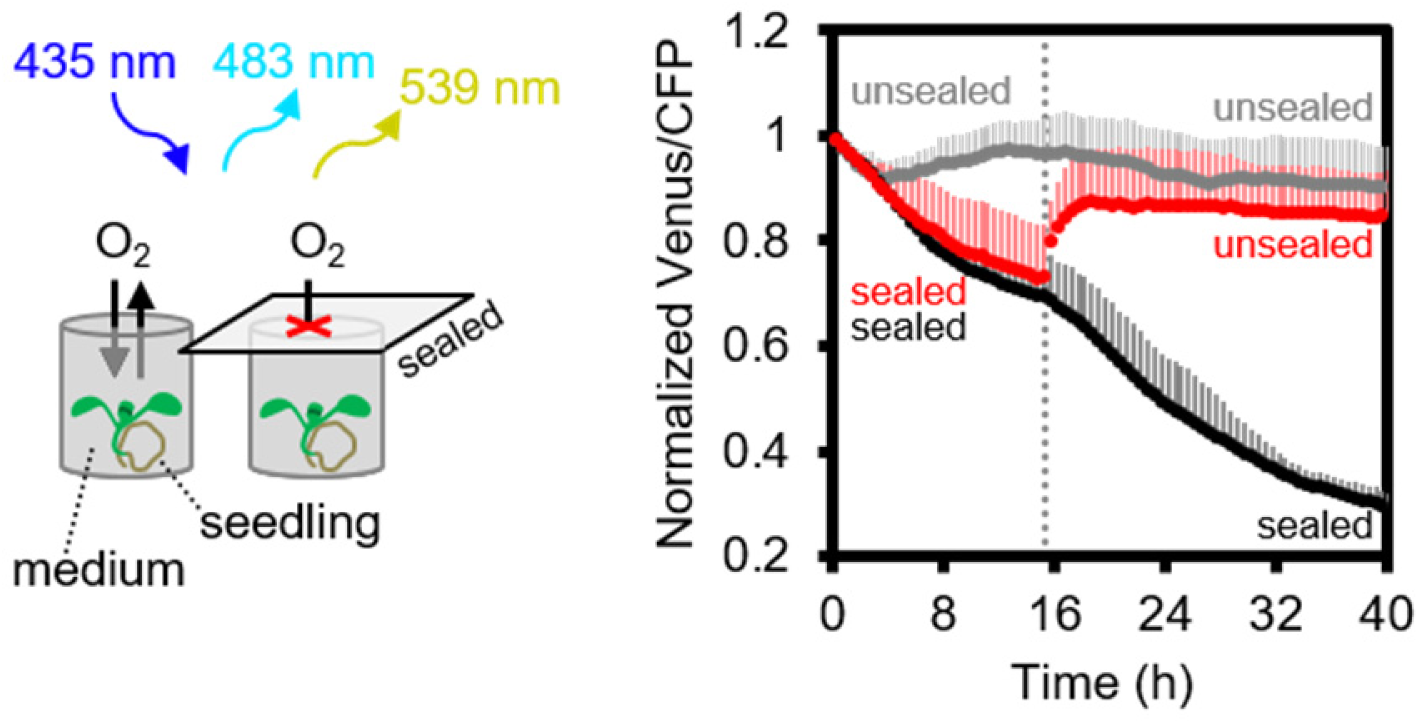
Arabidopsis MgATP^2−^ dynamics under low oxygen. Ten-day-old Arabidopsis seedlings, grown vertically on plates, were submerged in imaging medium on 96-well microtiter plates. Per well, three seedlings expressing no sensor (Col-0) or cytosolic ATeam were excited at 435 ± 10 nm and the emission at 483 ± 9 nm (mseCFP) and 539 ± 6.5 nm (cp173-Venus) was recorded. Wells were either left open (grey), sealed with an oxygen-proof, transparent qPCR film (black) or sealed for 15.5 h before the film was removed to reoxygenate the samples (red). Emission in wells with Col-0 plants was averaged and subtracted from that of ATeam-expressing plants to correct for background fluorescence. Data shown and used for background subtraction is the mean of 9-12 wells and error bars are SD.

## Discussion

### Fluorescent monitoring to understand ATP dynamics

Fluorescent ATP sensors were initially engineered for mammalian cells and tissues (Berg et al., 2009; Imamura et al., 2009; Kotera et al., 2010; Nakano et al., 2011; Tantama et al., 2013), where their use has allowed novel insights, such as subcellular ATP concentration gradients between compartments *in situ*, visualisation of stimulus-induced energy dynamics in neurons, responsiveness of ATP-sensitive K^+^ channel activity in single cells and synchronisation of Ca^2+^ and ATP dynamics in HeLa cells at histamine stimulation. With the exception of one report on cell death induced by hypersensitive response (Hatsugai et al., 2012), fluorescence ATP measurements have been lacking in plant research. Live-cell fluorescent monitoring complements widespread standard approaches, such as luminescent or HPLC-based ATP determinations in biological extracts, or radioisotope-based techniques for membrane transport assays (Manfredi et al., 2002; Khlyntseva et al., 2009; Lorenz et al., 2015; Monne et al., 2015). While those techniques offer high sensitivity and accuracy in quantifying ATP, and other nucleotides, in extracts, they provide endpoint measurements after removal from the functional biological system and have limited use for resolving the ATP concentration over time at the (sub)cellular and tissue level. Yet, high flux rates, rapid fluctuations and (sub)cellular gradients are fundamental characteristics of cellular energy physiology. Non-destructive live measurement of ATP has been available by ^31^P-NMR spectroscopy, which offers the additional benefit of also measuring ADP and other di- and tri-nucleotides (Gout et al., 2014). However, ^31^P-NMR is limited in spatial resolution and sensitivity. The superior sensitivity of fluorescence-based MgATP^2−^ monitoring is demonstrated in the experiments with isolated Arabidopsis mitochondria (Figure 3), where changes in the MgATP^2−^ concentration were monitored with time resolution in the order of seconds on only 20 μg mitochondrial protein. The spatial resolution of fluorescent ATP sensing is highlighted by its ability to detect MgATP^2−^ levels in individual cells (Figure 7) and single chloroplasts (Figure 1; ; Figure 5 – figure supplement 2), making ATP-binding probes a particularly versatile technique in tissue and cell physiology.

### Arabidopsis lines for monitoring MgATP^2−^ in the cytosol, chloroplasts and mitochondria

Governance of cytosolic ATP levels by two bioenergetic organelles in green plant cells raises pressing questions about the regulatory basis of subcellular ATP control (Gardeström and Igamberdiev, 2016). The cytosolic sensor lines indicated no differences in ratio between cytosol and nucleoplasm, indicative of rapid diffusion between the two locations and in line with previous observations for free Ca^2+^ concentrations and glutathione redox potential (Loro et al., 2012; Schwarzländer et al., 2016). The chloroplast sensor lines provide a good FRET signal, even in the presence of chlorophyll (Figure 5 – figure supplement 2). These lines will be a valuable tool to dissect the impact of photosynthesis on subcellular ATP, and investigations are currently underway in our labs. While no developmental phenotype was observed for the cytosolic and plastidic sensor lines (Figure 1; Figure 1 – figure supplements 2, 3), the strong phenotype that systematically resulted from sensor targeted to the mitochondrial matrix (Figure 1C; Figure 1 – figure supplement 2) requires caution. It cannot be ruled out that ATP homeostasis is generally perturbed in those sensor lines, although sensor-based measurements in mammalian cells have also indicated lowered ATP in the matrix, which can be accounted for by membrane potential-driven AAC-mediated ATP export (Imamura et al., 2009). The reason for the stunted phenotype is currently unclear. Interference with matrix ATP homeostasis by ATP buffering appears unlikely, especially since similar phenotypes of variable severity can also be observed for other mitochondrially targeted sensors (Figure 1 – figure supplement 4); similar issues were also noted in yeast (Schwarzländer et al., 2016). Clogging of the TIM/TOM machinery by import arrest of the synthetic sensor construct may account for the observed effects.

### Sensor characteristics and limitations revealed in vitro

Purified ATeam1.03-nD/nA was selective for MgATP^2−^ (Figure 2), refining previous characterisations of the parent sensor ATeam1.03 (Imamura et al., 2009). The sensor does not report on energy charge set by the ATP:ADP ratio directly (Pradet and Raymond, 1983), without an additional assumption that the total adenosine pool is constant. *In vivo, de novo* synthesis and degradation are unlikely to change the total adenosine pool as rapidly as ATP/ADP/AMP-cycling by (de-)phosphorylation. Hence, the MgATP^2−^ changes induced by inhibitor treatments and during hypoxia are likely to also reflect decreased energy charge. By contrast, differences in steady-state ATP levels between cells and tissues and organs may reflect constitutive differences in total adenylate pool sizes. Combinatorial analyses with sensors that directly respond to energy charge, such as variants of Perceval (Tantama et al., 2013), will be desirable in the future. However, the currently available ATP:ADP sensors suffer from serious pH sensitivity, which adds a level of complexity to the meaningful interpretation *in planta*, where pronounced pH fluctuations can occur.

### A dynamic assay to dissect ATP fluxes of isolated mitochondria

A key innovation from fluorescent protein-based sensing has been the ability to monitor biochemistry as it occurs in the intact biological system. Nevertheless, mechanistic conclusions have often been problematic based on attempts to link *in vitro* reductionism with *in vivo* complexity, and the challenges in bringing intact systems under experimental control. The ATP assays that we establish for isolated mitochondria introduce a useful intermediate (Figures 3 and 4). Despite the use of whole organelles as complex multi-functional units, the different steps of ATP dynamics can be monitored and interpreted mechanistically. Substrate feeding and inhibitor treatments allow for tight control over the functional state of the organelle, and controlled manipulation of ATP physiology. For example, we were able to distinguish the phosphorylation activities of ATP synthase and AK, and monitor either activity by exploiting their differential characteristics, including their sub-mitochondrial exposure to the matrix and the intermembrane space, respectively. Perturbation of mitochondrial function was observed with high sensitivity and kinetically resolved following inhibition of defined players in mitochondrial ATP production. Substrates and inhibitors are well suited to modulate respiratory activity, capacity and efficiency, and to induce transitions in ATP dynamics. They may be applied to dissect bioenergetic rearrangements in respiratory mutants in the future.

The current understanding of cellular bioenergetics has been driven by dynamic measurements in isolated mitochondria and chloroplasts, e.g. of oxygen consumption or Ca^2+^ transport. Yet, monitoring ATP dynamics, as the central product, has typically relied on indirect inference, or reconstruction of time-resolved data from individual samples (Attucci et al., 1991). The fluorescent biosensor-based ATP assay concept introduces a complementary method to continuously and dynamically monitor ATP transport fluxes from and into cells and isolated organelles, as demonstrated using plant mitochondria. Since the sensor protein is added externally to the samples, and does not require transgenic sensor expression by the samples, cell components from any species can be studied for ATP generation, consumption and membrane transport. The assays may be further expanded to study membrane transport dynamics. For instance, the response of ATP transport across the chloroplast envelope to changes in photosynthetic activity will be readily detectable making it possible to address yet unresolved questions about transport efficiency, directionality and modes, under controlled external conditions (Haferkamp et al., 2011; Gardeström and Igamberdiev, 2016). The assay principle is adaptable to other subcellular structures, and potentially even liposome, cell, and tissue systems. It may also facilitate the improvement of ATP-coupled enzyme assays. Monitoring ATP export and uptake by cells may be of specific interest to address current questions about intercellular ATP fluxes and the signalling function of extracellular ATP (Kim et al., 2006; Tanaka et al., 2010; Choi et al., 2014a). ATeam variants with higher affinities to match lower ATP concentration ranges have been engineered, but are likely MgATP^2−^ specific (Imamura et al., 2009).

### Monitoring ATP in planta by three sensor detection approaches

We have performed live sensing of ATP *in planta* by three distinct *in vivo* techniques. Microtiter plate-based fluorimetry (Figures 3-5, and 10), CLSM (Figures 1, 6, and 7) and LSFM (Figures 8 and 9) complement another to investigate the dynamics and distribution of ATP in living plant tissues. While CLSM has been extensively used in combination with other fluorescent protein sensors in plants, LSFM has only recently been adjusted to enable FRET-based Ca^2+^ measurements in Arabidopsis roots (Costa et al., 2013; Candeo et al., 2017). Microtiter plate-based fluorimetry has been widely used for cultured animal cells and yeast (Birk et al., 2013; Morgan et al., 2016), but rarely in plant tissues (Rosenwasser et al., 2010; Rosenwasser et al., 2011), and a rigorous technical validation was missing. Our analysis shows that careful adjustment of fluorimetric settings, with the sensor properties and the characteristics of the tissue can make plate reader-based measurements a robust approach for continuous monitoring of fluorescent protein sensors over extended time periods. While heterogeneity between cells and tissues is averaged, large sample numbers can be assessed in parallel enabling robust controls and accounting for biological variability. Genetic targeting of the sensor to a specific subcellular location where it responds to the local ATP levels means that the resolution of subcellular physiology is maintained. Further, side-by-side measurements of different sensors in different tissues or cell compartments may allow monitoring and dissecting the interplay of different physiological responses to the same stimulus. For instance, the impact of hypoxia on cyto-nuclear ATP, pH and Ca^2+^, or the impact of illumination on the ATP concentration in the cytosol and the chloroplast stroma may be assessed. Parallel side-by-side monitoring provides an alternative to multiplexing of several sensors co-expressed in the same plant. The robustness of the assay may even be suitable for genetic or chemical biological screens (Dejonghe and Russinova, 2017) based on fluorescent sensors in the future.

### Sensing and manipulating MgATP^2−^ in vivo

Decreasing and increasing the cytosolic MgATP^2−^ levels *in vivo* by chemical treatments resulted in a reliable response that covered nearly the full theoretical dynamic range of the sensor (Figure 5). We infer that the sensor is functional *in vivo*, and not affected by proteolytic cleavage. The effective decrease of MgATP^2−^ after inhibition with antimycin A validated the role of mitochondria as major suppliers of cytosolic ATP, as the loss of respiration-derived ATP could not be replaced. Nevertheless, the response kinetics to the chemical treatments appeared to be dominated by tissue uptake rate. Monitoring the treatments by CLSM indicated both rapid and slower responses depending on tissue type (Figure 6 – figure supplement 2). Yet, all tissues gradually adopted a similarly low FRET ratio of the MgATP^2−^ free sensor. The ability to drive the sensor to its extremes *in vivo* allows the conversion of *in vivo* FRET ratios to absolute MgATP^2−^ concentrations, as routinely done in Ca^2+^ sensing (Palmer and Tsien, 2006), and thiol redox sensing (Schwarzländer et al., 2008). Since the conversion requires additional assumptions, and becomes inaccurate towards the non-linear response range of the sensor, a fully quantitative approach has been criticized. In this work we use the FRET values as direct representation of the *in vivo* dynamics and relative differences (Wagner et al., 2015b). When conversion to absolute concentrations is desired, they should be regarded as an estimate. Here about 2 mM MgATP^2−^ can be estimated for the cytosolic ATP concentrations at steady state in the dark (Figures 2 and 5). This concentration value complements ATP determinations normalized to chlorophyll contents by protoplast fractionation (Stitt et al., 1982; Gardeström and Wigge, 1988) and by NMR (Gout et al., 2014).

### Towards an integrated understanding of energy physiology in development and under stress

CLSM resolved organ, tissue and cell differences in a detailed MgATP^2−^ map of the Arabidopsis seedling (Figures 6 and 7). Similar maps have been generated for hormone distribution (e.g. auxin and abscisic acid Brunoud et al., 2012; Jones et al., 2014) to study plant development. For central metabolites high-resolution transcriptional analysis has aimed to generate tissue maps (Chaudhuri et al., 2008), but direct *in vivo* mapping across tissues and organs has not been undertaken. Differences in the MgATP^2−^ concentration between tissues and cells have not previously been distinguishable with any reasonable resolution, and the MgATP^2−^ map therefore unveils a surprisingly heterogeneous and dynamic picture. Green tissues showed overall high MgATP^2−^ concentration, which are unlikely due to photosynthetic activity because the seedlings were dark-adapted. Etiolation did not only decrease MgATP^2−^ in the green tissues, but also increased levels in the roots. The biological significance of tissue specific MgATP^2−^ heterogeneity deserves in-depth dissection in the context of metabolism and development in the future. Since the map represents MgATP^2−^, as the main bioavailable form of ATP, differences may, in principle, be due to ATP pool size, pH and/or Mg^2+^. Many proteins bind MgATP^2−^, but the extent to which this affects the buffering of the pool and the free concentration, which is available for binding by the sensor, is currently unknown.

With the appropriate caution, the map provides insight into MgATP^2−^ distribution at system level and the concept sets a reference point for expansion into several dimensions, including time, subcellular compartment and other genetic, biochemical or physiological parameters. Time-lapse imaging will allow resolving the dynamic changes in a tissue, both during developmental processes and in response to external stimuli. Our observation that the MgATP^2−^ concentration increases with decreasing root hair growth exemplifies the occurrence of such dynamics and provides evidence for the complex relationship between cellular energy status and growth. Analogous maps can be generated for other subcellular compartments, such as the plastids, to obtain a tissue-resolved map of subcellular MgATP^2−^ heterogeneity. Other sensors, e.g. for pH, Ca^2+^ or glutathione redox potential, may be used and superimposed to gradually build up a comprehensive representation of the cell physiological status of the whole plant. Superimposition with transcriptomic and proteomic maps (e.g., Chaudhuri et al., 2008; Li et al., 2016) may even allow correlation across the organisational levels of the cell. Such systemic multi-dimension *in vivo* mapping will provide a novel foundation to modelling attempts of whole plant metabolism and to the understanding of how metabolism underpins plant development.

Disruption of fresh oxygen supply to Arabidopsis seedlings resulted in the rapid onset of a gradual decrease in the cytosolic MgATP^2−^ concentration (Figure 6), following expectations from previous work on plant extracts and using NMR (Xia and Saglio, 1992; Geigenberger et al., 2000; Gout et al., 2001; van Dongen et al., 2003). The decrease occurred in four phases and was fully reversible by re-oxygenation. FRET ratios initially declined slowly, however, after extended block of oxygen supply, a sharp decrease in the MgATP^2−^ concentration occurred to a new plateau level, which may represent the lower sensitivity limit of the probe. The exact kinetics depended on the experimental setup and the type of tissue, yet the overall response was reproducible (Figure 10; Figure 10 – figure supplement 1). The rapid response and the gradual decrease together support the rationales that the cytosolic MgATP^2−^ pool strictly depends on oxidative phosphorylation, that AK-based buffering may at most delay MgATP^2−^ depletion and that sharp MgATP^2−^ depletion can be avoided, probably by restructuring of metabolic fluxes (Geigenberger, 2003; Zabalza et al., 2009). The experimental setup did not generate a sudden decrease in oxygen availability, but relied on dark respiration for gradual oxygen depletion. The observed responses therefore represent the integrated effects of a dynamic progression of hypoxia and the dynamic metabolic changes; as they may occur in a situation of sudden (deep) water logging. Hypoxia-associated intracellular acidification may also impact on the sensor response. Yet, pH-induced changes are not artefacts and carry physiological meaning, since a decrease in the concentration of MgATP^2−^ does not only affect the sensor, but also the endogenous MgATP^2−^-dependent proteins. Further, the pH controlled MgATP^2−^ depletion treatments indicate only a minor contribution (Figure 5 – figure supplement 3) and prior work has suggested that cytosolic pH hardly decreases below 7, even under anoxia (Gout et al., 2001; Schulte et al., 2006). As such, the hypoxia assays demonstrate that ATeam1.03-nD/nA allows continuous monitoring of subcellular MgATP^2−^ pools in response to stress. Since the exact ATP kinetics are shaped by the cellular stress response machinery, they will provide a sensitive and integrated readout for defects or modifications in mutants.

### Conclusions and outlook

We have investigated mitochondrial bioenergetics, plant hypoxia responses and MgATP^2−^ content in plant tissues depending on growth conditions and development. These studies exemplify the versatility of fluorescent ATP sensing to open new doors in plant biology. Their systematic follow up and extrapolation will be required for a systems view of ATP in the future. Although ATP biochemistry has been extensively studied in the last century, surprisingly large gaps remain in our understanding of its dynamics within cells and whole plants. Sensing MgATP^2−^ dynamically and with (sub)cellular resolution adds novel depth to the study of plant metabolism, development, signalling and stress responses. Our understanding of plant-microbe interactions, where the biochemistry underpinning the localized and dynamic responses have been notoriously hard to capture, may benefit in particular. Other potential applications include the identification and *in vivo* validation of ATP transport systems, a better understanding of the coordination between plastids and mitochondria in ATP production, an appraisal of the impact of uncoupling systems, the visualisation of energy parasitism in diatoms, the extension of *in situ* enzyme assays and the mapping of MgATP^2−^ to other sensor outputs and oxygen gradients in tissues. Optimization of high affinity ATP sensors for the apoplast could support investigations on extracellular ATP signalling. Additional sensors for AMP, ADP and ATP:ADP ratio can be integrated into the methodological framework introduced here. Each time that a biosensor for a new facet of cell physiology, e.g. for free Ca^2+^ or glutathione redox potential (Allen et al., 1999; Pei et al., 2000; Meyer et al., 2007; Schwarzländer et al., 2008; Krebs et al., 2012; Loro et al., 2012; Wagner et al., 2015b), has been introduced into plant research, this has yielded a burst of discovery. We expect ATP sensing to be no exception.

## Materials and Methods

### Cloning of sensor constructs and generation of plant lines

The ATeam1.03-nD/nA sequence was PCR-amplified from pENTR1A:ATeam1.03-nD/nA. The leader sequence from *Nicotiana plumbaginifolia* β-ATPase (Logan and Leaver, 2000) for mitochondrial import was fused to the N-terminus by extension PCR. For constitutive plant expression under a CaMV 35*S* promoter, this fusion and the unfused sequence for cyto-nuclear targeting were subcloned into pDONR207 (Invitrogen Ltd) and ultimately pB7WG2 and pH2GW7, respectively (Karimi et al., 2002). For targeting to the plastid stroma, the leader sequence from *Nicotiana tabacum* transketolase (Wirtz and Hell, 2003; Schwarzländer et al., 2008) and the ATeam1.03-nD/nA sequence were subcloned into pENTR/D-TOPO (Invitrogen Ltd) via NdeI/PstI and BamHI/XbaI restriction sites, respectively. The fusion was cloned into pEarleyGate100 (Earley et al., 2006) for 35*S*-driven expression. Primer sequences are detailed in **Supplementary File 1**. The non-fused sequence was also inserted into pETG10A for protein expression *in Escherichia coli* cells. Agrobacterium-mediated transformation of *Arabidopsis thaliana* (L.) Heynh. (accession Columbia, Col-0) was performed by floral dip (Clough and Bent, 1998). Transformants and homozygous lines were selected by chemical resistance and fluorescence intensity. Generation of the NES-YC3.6 Cameleon line was described previously (Krebs et al., 2012).

### Chemicals

Chemicals were purchased from Sigma-Aldrich. Stock solutions of ATP, ADP and AMP were freshly supplemented with equimolar concentrations of MgCl2 except for Mg^2+^ titration. All stock solutions were adjusted to assay pH prior to use.

### Plant culture

Where not indicated otherwise, Arabidopsis seedlings were grown from surface-sterilized seeds on vertical plates containing half-strength Murashige and Skoog (MS) medium (Murashige and Skoog, 1962) with 1% (w/v) sucrose and 0.8% (w/v) Phytagel under long-day conditions (16 h 80-120 μmol photons m^−2^ s^−1^ at 22°C, 8 h dark at 18°C) after stratification at 4°C in the dark. For line preparation of leaf discs, plants were germinated and grown on soil under long-day conditions (17°C, 16 h 50-75 μmol photons m^−2^ s^−1^, 8 h dark) after stratification at 4°C in the dark.

### Plant phenotyping

Seeds were stratified at 4°C in the dark for 2 d on half-strength MS medium + 1 % (w/v) sucrose + 1 % (w/v) MES + 1 % (w/v) Phytagel, pH 5.8 and seedlings were grown on the plates vertically side by side with their corresponding controls for 5 d under long-day conditions (16 h at 22°C and 75-100 μmol photons m^−2^ s^−1^, 8 h at 18°C and darkness). Primary root length was documented and quantified using ImageJ. Plants were then individually transferred to Jiffy-pots, randomly distributed on standard greenhouse flats and grown in long-day (16 h at 19°C and 60-80 μE m^−2^ s^−1^, 8 h at 17°C and darkness) growth chambers. Leaf rosette development was documented photographically and rosette size was analysed with the custom Leaf Lab tool (Version 1.41) as described previously (Wagner et al., 2015b). Height of the primary inflorescence was systematically captured with a camera and quantified using ImageJ. Siliques were manually counted when the first siliques turned yellow but had not yet shattered.

### Purification of ATeam1.03-nD/nA

*E. coli* strain Rosetta 2 (DE3) carrying pETG10A-ATeam1.03-nD/nA was grown in Lysogeny Broth (LB; Bertani, 1951) medium at 37°C to an OD_600_ of 0.2. The culture was transferred to 20°C until cells reached an OD_600_ of 0.6-0.8. Expression of 6×His-ATeam1.03-nD/nA was induced by isopropyl β-D-1-thiogalactopyranoside at a final concentration of 0.2 mM overnight at 20°C. Cells were collected by centrifugation at 4,000 *g* for 10 min at 4°C and the pellet was resuspended in lysis buffer (100 mM Tris-HCl, pH 8.0, 200 mM NaCl, 10 mM imidazole) supplemented with 1 mg/mL lysozyme, 0.1 mg/mL DNaseI (Roche) and cOmplete protease inhibitor cocktail (Roche). After 30 min incubation on ice, cells were sonicated (3 × 2 min, 40% power output, 50% duty cycle). The lysate was centrifuged at 40,000 *g* for 40 min at 4°C and the supernatant was loaded onto a Ni-NTA HisTrap column (GE Healthcare) and proteins were eluted with an imidazole gradient (10-200 mM in 100 mM Tris-HCl, pH 8.0, 200 mM NaCl) using an ÄKTA Prime Plus chromatography system (GE Heathcare). Fractions containing ATeam1.03-nD/nA were pooled, concentrated by ultrafiltration and applied to a HiLoad 16/600 Superdex 200 column (GE Healthcare) pre-equilibrated with 20 mM Tris-HCl, pH 8.0, 150 mM NaCl. Fractions containing intact ATeam1.03-nD/nA were pooled, concentrated by ultrafiltration, supplemented with 20% (v/v) glycerol and stored at − 86°C.

### Characterization of ATeam1.03-nD/nA in vitro

Concentration of purified ATeam1.03-nD/nA was quantified according to (Bradford, 1976). Protein at a final concentration of 1 μM was mixed with basic incubation medium (0.3 M sucrose, 5 mM KH_2_PO_4_, 50 mM TES-KOH, pH 7.5, 10 mM NaCl, 2 mM MgSO_4_, 0.1% (w/v) BSA) for all *in vitro* assays, except for the Mg^2+^ titrations, in which MgSO_4_ was omitted from the basic incubation medium. A FP-8300 spectrofluorometer (Jasco) at 25°C was used to excite mseCFP at 435 ± 5 nm and emission spectra between 450 to 600 nm were recorded with a bandwidth of 5 nm. Venus/CFP ratios were calculated as measure of FRET efficiency from the fluorescence emission intensities at 527 nm (cp173-Venus) and 475 (mseCFP) nm after blank correction.

### Mitochondrial isolation and oxygen consumption assays

Arabidopsis seedlings were grown in hydroponic pots for 14 days as described by Sweetlove et al. (2007) under long-day conditions (16 h 50-75 μmol photons m^−2^s^−1^ at 22°C; 8 h dark at 18°C). Seedling mitochondria were isolated as described by (Sweetlove et al., 2007; Schwarzländer et al., 2011). Oxygen consumption was assayed as described by (Sweetlove et al., 2002; Schwarzländer et al., 2009) using two Clark-type electrodes (Oxytherm, Hansatech).

### Multiwell plate reader-based fluorimetry

ATeam1.03-nD/nA was excited with monochromatic light at a wavelength of 435 ± 10 nm in a CLARIOstar plate reader (BMG Labtech). Emission was recorded at 483 ± 9 nm (mseCFP) and 539 ± 6.5 nm (cp173-mVenus) using transparent 96-well plates (Sarstedt). The internal temperature was kept at 25°C and the plate was orbitally shaken at 400 rpm for 10 s after each cycle. For assays with isolated mitochondria, 20 μg total protein in basic incubation medium (0.3 M sucrose, 5 mM KH_2_PO_4_, 50 mM TES-KOH, pH 7.5, 10 mM NaCl, 2 mM MgSO_4_, 0.1% (w/v) BSA) were supplemented with 1 μM purified ATeam1.03-nD/nA in a total volume of 200 μL per well. Fluorescent background of the basic incubation medium was recorded and subtracted from all data before analysis. For *in vivo* experiments with Arabidopsis seedlings and leaf discs, plant material was submerged in 10 mM MES, pH 5.8, 5 mM KCl, 10 mM MgCl_2_, 10 mM CaCl_2_. Plates were kept in the dark for at least 30 min before recording to minimize potential effects of active photosynthesis. TES buffer was replaced with Bis-Tris-HCl (covering pH values between 6.0 and 7.0) or Tris-HCl (covering pH values between 7.5 and 8.5) where indicated. To restrict supply with oxygen, plates were sealed with ultra-clear films for qPCR (VWR).

### Confocal laser scanning microscopy and analysis

Confocal imaging was performed at 20°C using a Zeiss LSM780 microscope and a ×5 (EC Plan-Neofluar, 0.16 N.A.), ×10 (Plan-Apochromat, 0.3 N.A) or ×40 lens (C-Apochromat, 1.20 N.A., water immersion) using the procedure described previously (Wagner et al., 2015a). ATeam1.03-nD/nA was excited at 458 nm and fluorescence of mseCFP and cp173- mVenus were measured at 465-500 nm and 526-561 nm, respectively, with the pinhole set to 3 airy units. Chlorophyll fluorescence was collected at 650-695 nm. MitoTracker Orange was excited at 543 nm and emission was recorded at 570-619 nm. Plants were dark-adapted for at least 30 min before image acquisition. Single plane images were processed with a custom MATLAB-based software (Fricker, 2016) using *x*,*y* noise filtering and fluorescence background subtraction.

### Light sheet fluorescence microscopy and analysis

Hair cell growth on roots from 6-day-old seedlings was followed for 10 min with images acquired every 3 s. Each image was generated as a maximum intensity projection (MIP) of 15 stacks spaced by 3 μm and the MIP acquisition time was 1 s, based on 50 ms exposure per stack and data transfer time. The LSFM used was specifically designed to study plant roots (Costa et al., 2013; Candeo et al., 2017). The source was a single-mode fibre-coupled laser emitting radiation at 442 nm (MDL-III-442, CNI) to excite ATeam mseCFP. A cylindrical lens in combination with a 10× water-dipping microscope objective (Olympus UMPLFLN 10×W, 0.3 N.A.) created a thin sheet of laser light on the sample (5-μm-thick and 800-μm-high in the vertical direction). The detection unit consisted of a 20× water-dipping microscope objective (Olympus UMPLFLN 20×W, 0.5 N.A.), held orthogonally to the excitation axis. An image splitter (dichroic filter at 505 nm, DMLP505, Thorlabs; band-pass filters, MF479-40 and MF535-22, Thorlabs) and a fast camera (Neo 5.5 sCMOS, ANDOR) enabled the simultaneous wide-field acquisition of two spectrally different images, as required for a FRET indicator. The system provided an intrinsic optical sectioning with minimal light exposure of the sample, permitting a fast volumetric acquisition and long-term imaging over a wide-field of view with single-cell detail. The seedlings were grown vertically in fluorinated ethylene propylene tubes filled with half-strength MS medium, 0.1% (w/v) sucrose; 0.05% (w/v) MES, pH 5.8 (Tris), solidified by 0.5 % (w/v) Phytagel as described in (Candeo et al., 2017) and kept in an imaging chamber filled with liquid MS-based medium lacking sucrose (half-strength MS medium; 0.05% (w/v) MES; pH 5.8 (Tris)).

The Venus/CFP ratio at the tip of single root hairs was extracted with FIJI (https://fiji.sc/) after image registration by using the Fiji plugin “Template Matching” (https://sites.google.com/site/qingzongtseng/template-matching-ij-plugin). Finally, the Matlab Fast Fourier Transform was used to calculate the Power Spectral Densities from the normalized Venus/CFP time courses (Candeo et al., 2017). To quantify the average speed growth of root hairs we measured their elongation during the entire acquisition and divided by the time (10 min).

## Acknowledgements

We thank Takeharu Nagai (Osaka, Japan) for the kind gift of the pENTR1A:ATeam1.03- nD/nA construct, Jan Riemer (Cologne, Germany), Elisa Petrussa and Valentino Casolo (Udine, Italy) for important discussions, and Silvina Paola Denita Juárez (Buenos Aires, Argentina) and Andrea Magni (Milan, Italy) for assistance with the LSFM experiments.

## Author contributions

V.D.C. performed *in vitro* experiments, analysed the data, and assisted in writing the article. P.F., T.N and M.E assisted in optimizing and performing the *in vivo* measurements and analysed data. V.C.P. generated the chloroplast sensor lines. A.Ca. optimized LSFM, provided data processing protocols and analysed data. I.S. performed cloning and preliminary *in vitro* characterisations. M.D.F. contributed custom imaging analysis software, and consulted on the data analysis. C.G. contributed to the quantitative FRET analyses and assisted in writing the article. I.M.M. contributed to the data analysis and interpretation, and assisted in writing the article. A.B. built the LSFM, provided measurement routines and advice on data collection. B.L.L. and M.Z. co-supervised research, and assisted in writing the article. A.J.M. provided imaging expertise and microscopy facilities. A.Co. co-supervised the research, designed and performed LSFM experiments and assisted in writing the article. S.W and M.S. jointly conceived, performed and supervised the research, analysed the data and wrote the article.

## Figures

**Figure 1 - figure supplement 1:**
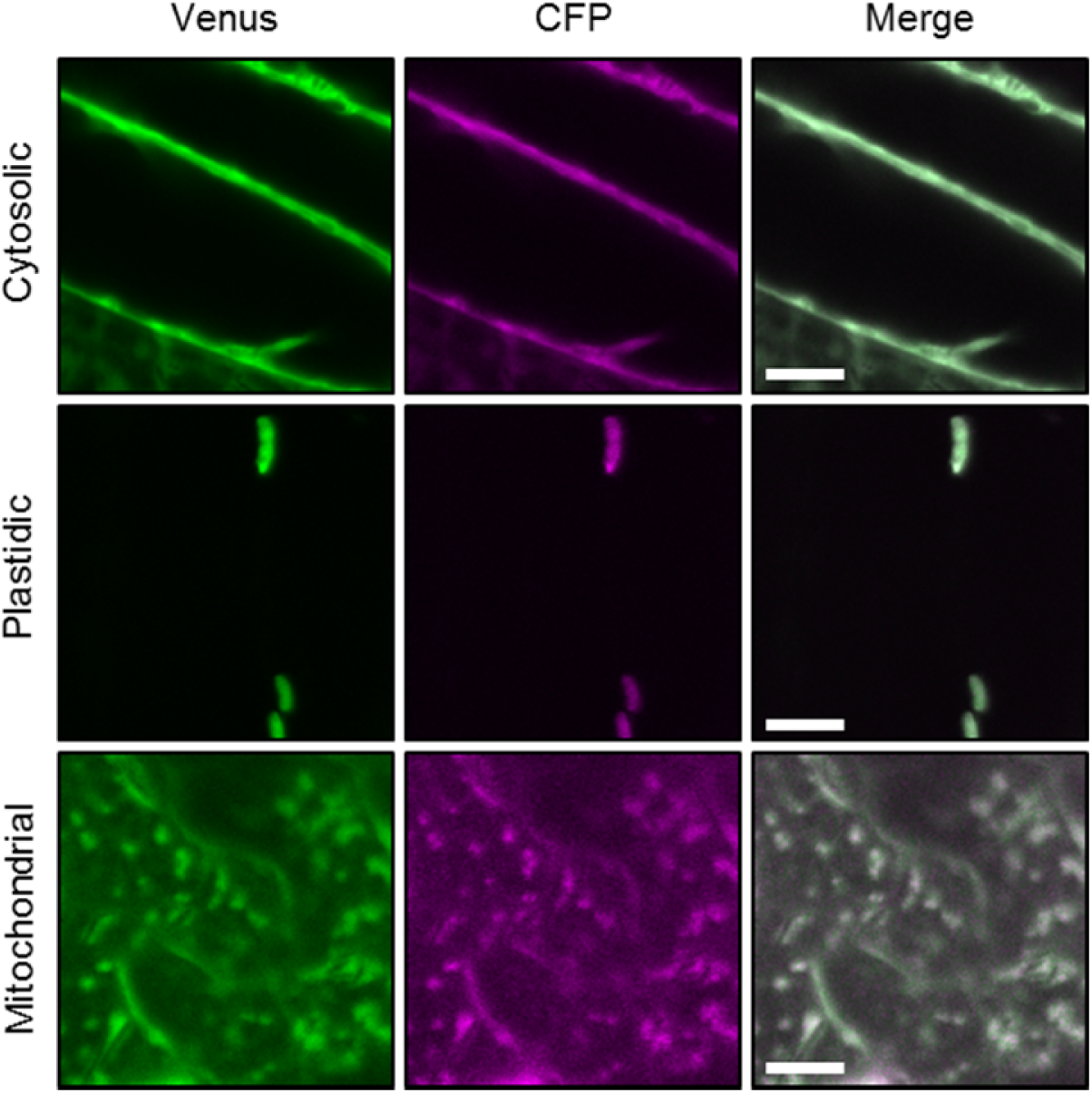
Ratiometric imaging of ATeam in cellular compartments of Arabidopsis seedlings. Five-day-old seedlings grown vertically on half-strength MS + 1% (w/v) sucrose were imaged by CLSM. Fluorescence of Venus (green) and CFP (magenta) was assessed in hypocotyl cells using the same settings except for power of the 458 nm laser which was set to 10 % of maximal power for the nuclear-cytosolic and plastidic and 30 % for the mitochondrial line. Scale bar = 10 μm. The ratiometric analysis of the images is shown in Figure 1C.

**Figure 1 – figure supplement 2:**
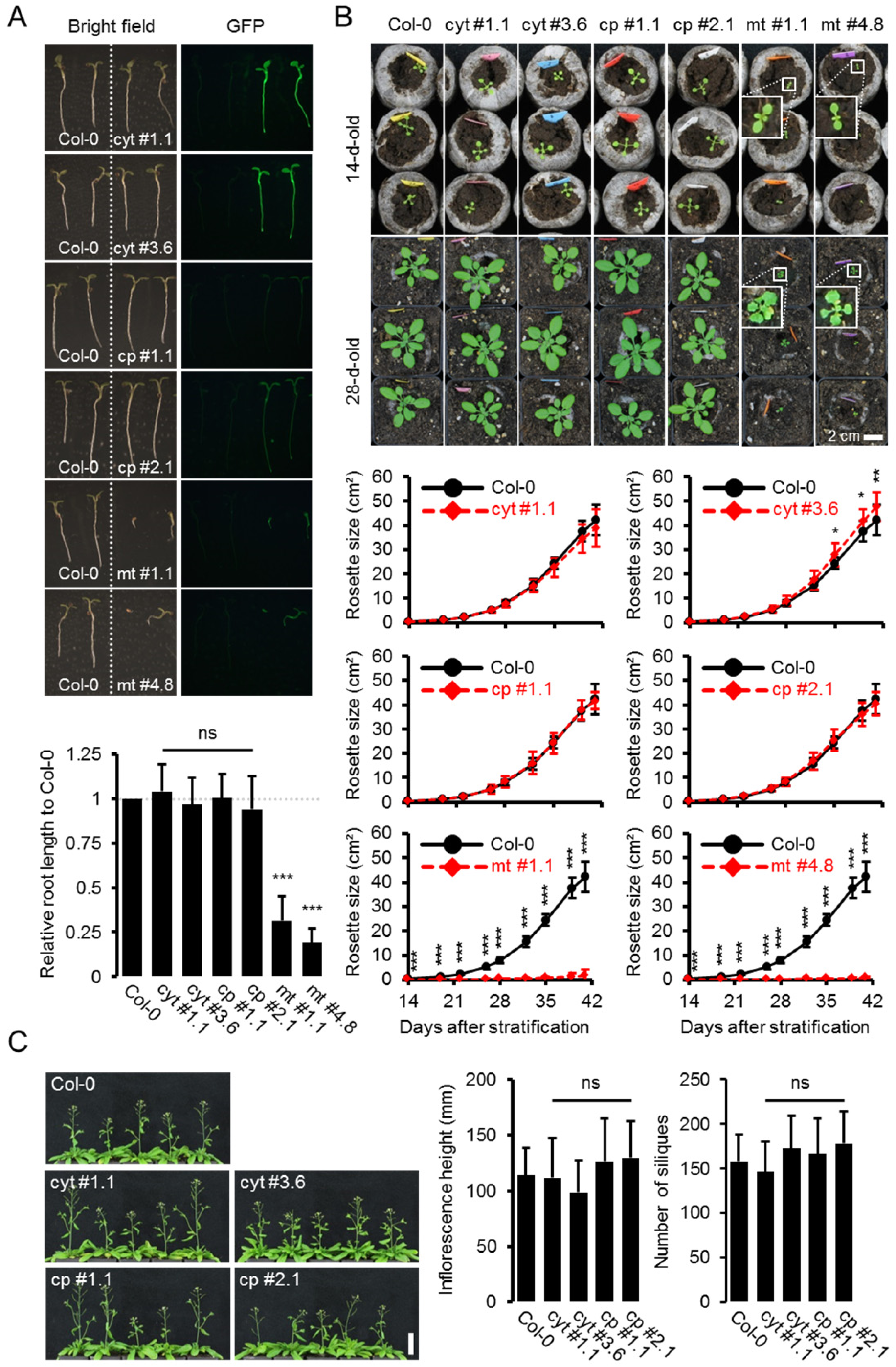
Whole plant phenotyping of homozygous ATeam lines. (A) Plants homozygously expressing ATeam in the cytosol (cyt), plastids (cp) or mitochondria (mt) were grown vertically for 5 days on solidified half-strength MS medium side by side with wild-type Col-0. Fluorescence of plant material was checked with an epifluorescence microscope equipped with a GFP filter. Primary root length of 35 ATeam and 35 Col-0 plants per independent sensor line was quantified 5 days after stratification and statistical difference from Col-0 was tested with a one-way ANOVA followed by the Dunnett test. ns: p > 0.05, ***p ≤ 0.001; error bars = SD. (B) 20 (Col-0, cyt, cp) and 12 (mt) randomly selected plants shown in (A) were transferred to soil and grown in long-day growth chambers. Rosette development was documented photographically and the leaf rosette area was analysed until the majority of plants developed first open flowers. Growth curves for each sensor line are plotted against the same set of Col-0 plants and statistical differences were assessed for individual timepoints separately with a one-way ANOVA, followed by the Dunnett test. *p ≤ 0.05, ***p ≤ 0.001; error bars = SD. (C) Primary inflorescence height of 46-d-old plants was captured photographically and quantified. Siliques of 61-d-old plants were manually counted. *n* = 20; error bars = SD; ns: p > 0.05 (one-way ANOVA with Dunnett test to compare sensor lines with Col-0). Scale bar = 50 mm. Plants expressing mitochondrial ATeam did not bolt until day 61 and were therefore not included in the analysis.

**Figure 1 – figure supplement 3:**
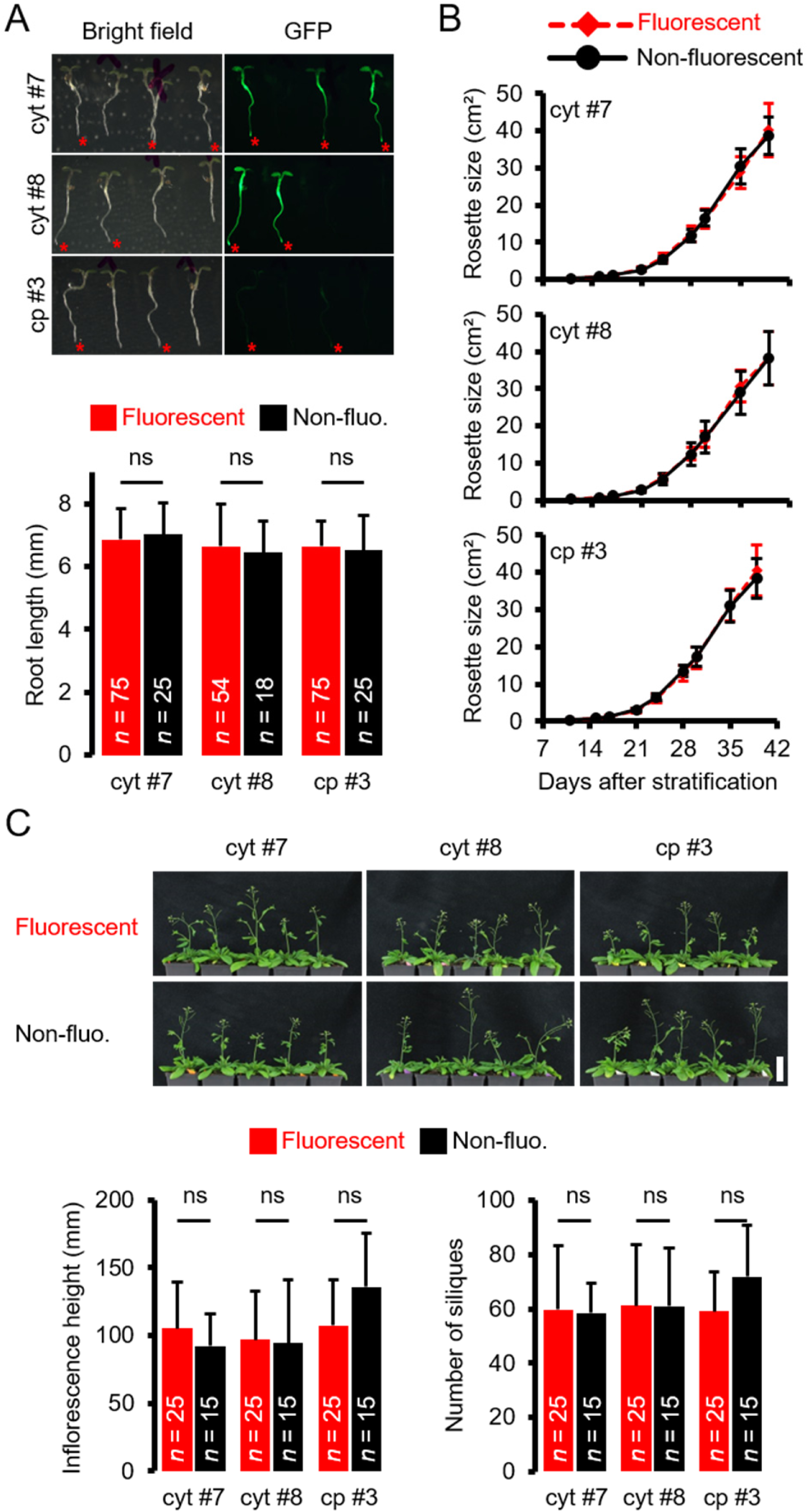
Whole plant phenotyping of heterozygous ATeam lines. (A) Plants heterozygously expressing ATeam in the cytosol (cyt) or plastids (cp) were grown vertically for 5 days on solidified half-strength MS medium. Fluorescence of plant material was checked with an epifluorescence microscope equipped with a GFP filter. Fluorescent individuals are marked by a red asterisk. Primary root length of fluorescent and non-fluorescent plants in a 3:1 ratio was quantified 4 days after stratification and statistical difference was tested with a one-way ANOVA followed by the Tukey test. ns: p > 0.05; error bars = SD. (B) 25 fluorescent and 15 non-fluorescent plants per independent line were randomly selected, transferred to soil and grown in long-day growth chambers. Rosette development was documented photographically and the leaf rosette area was analysed until the majority of plants developed first open flowers. Growth curves of fluorescent and non-fluorescent individual are plotted for each line and no statistical differences at any time point were found with a one-way ANOVA, followed by the Tukey test. (C) Primary inflorescence height of 43-d-old plants was captured photographically and quantified. Siliques of 56-d-old plants were manually counted. ns: p > 0.05 (one-way ANOVA with Tukey test); error bars = SD. Scale bar = 50 mm. Plants expressing mitochondrial ATeam did not bolt until day 56 and were therefore not included in the analysis.

**Figure 1 – figure supplement 4:**
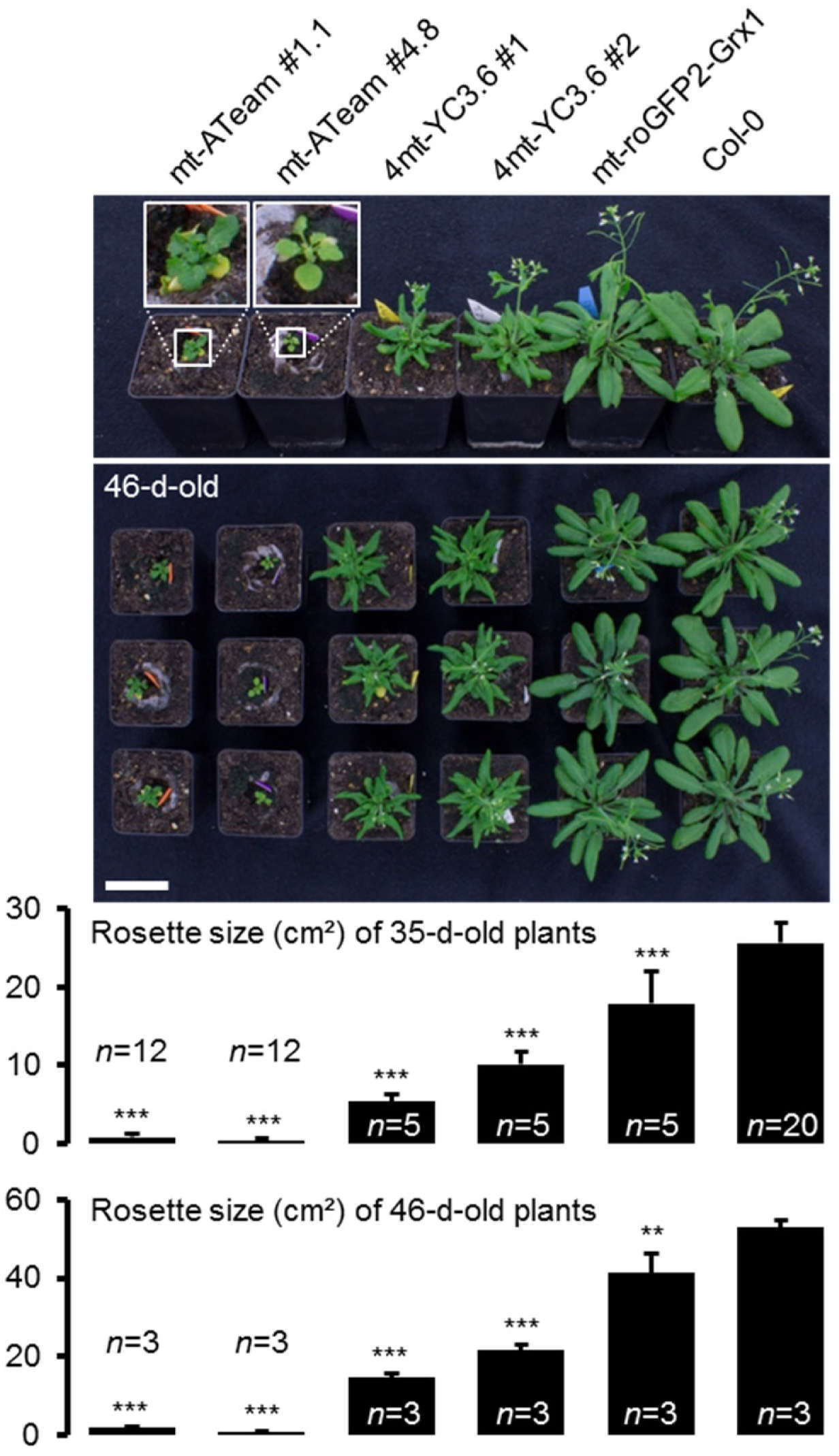
Phenotyping of Arabidopsis expressing mitochondrial sensor proteins. Plants homozygously expressing mitochondrial ATeam, YC3.6 and roGFP2-Grx1 (*n* = 5 for each independent line) constitutively under a CaMV *35S* promoter were grown vertically for 5 days on solidified half-strength MS medium, transferred to soil and grown under long-day conditions. Total rosette size was quantified 35 days after stratification and compared with the same set of wild-type Col-0 plants as shown in Fig. 1 – figure supplement 2 (*n* = 20). Three representative plants of each line were selected for the shown lower photograph 46 days after stratification which was additionally used to quantify the rosette area (lower graph). Statistical difference from Col-0 was tested with a one-way ANOVA followed by the Dunnett test. **p ≤ 0.01, ***p ≤ 0.001; error bars = SD. Scale bar = 5 cm.

**Figure 2 - figure supplement 1:**
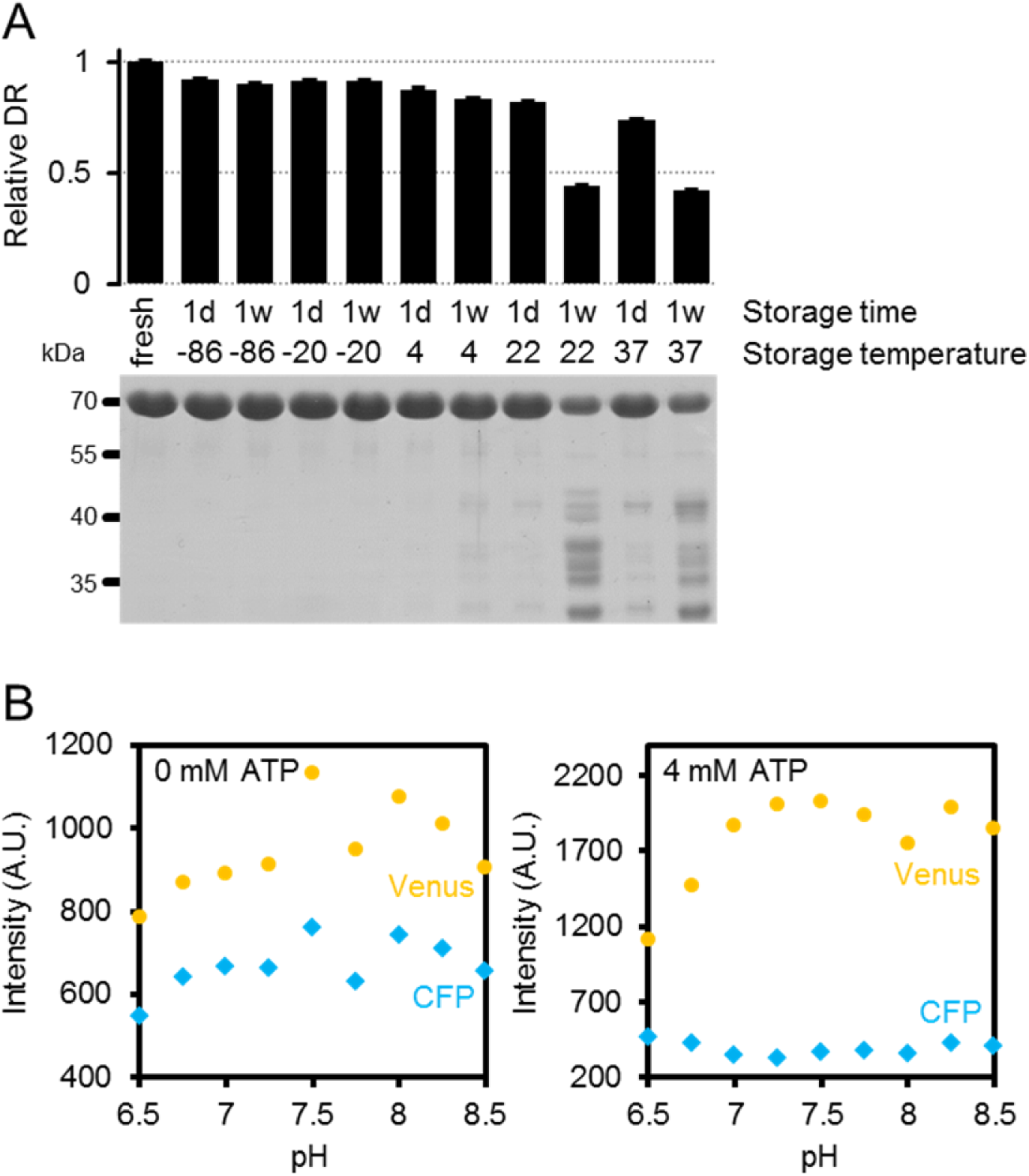
Characteristics of purified ATeam1.03-nD/nA. (A) Freshly purified ATeam was stored for one day (1d) or one week (1w) at temperatures between −86 and 37°C. The dynamic range (DR) of stored protein represents the ratio of Venus/CFP at zero and saturating ATP (4 mM in the presence of Mg^2+^ in excess) and was normalized to freshly purified protein. Protein degradation in these samples was analysed by SDS-PAGE. The expected molecular mass of intact 6×His-ATeam1.03-nD/nA is 70.1 kDa. (B) Fluorescence emission intensity (excitation at 435 ± 5 nm) of the individual fluorescent proteins mseCFP and cp173-Venus at variable pH and 0 or 4 (saturating) mM ATP. Data is averaged from four technical replicates and error bars represent SD but are too small to be visibly displayed.

**Figure 5 - figure supplement 1:**
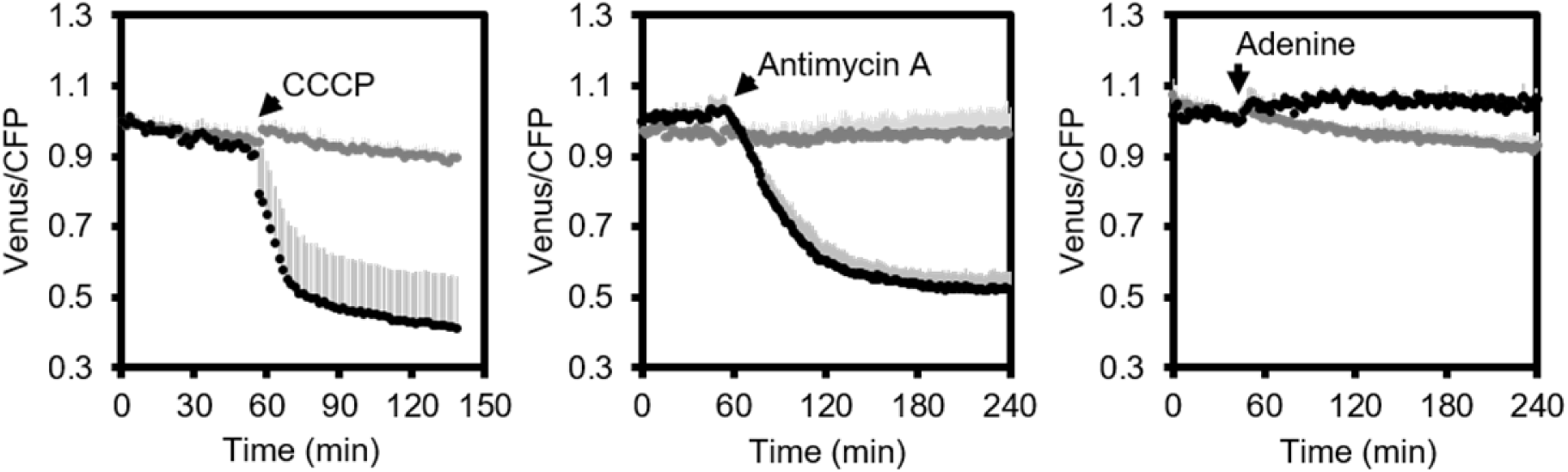
Fluorometric readings of MgATP^2−^ dynamics in Arabidopsis seedlings. Independent repetition of experiments shown in Figure 5C-E. Per well, two 7-day-old Arabidopsis seedlings expressing no sensor (Col-0) or cytosolic ATeam were excited at 435 ± 10 nm and the emission at 483 ± 9 nm (mseCFP) and 539 ± 6.5 nm (cp173-Venus) was recorded. CCCP (100 μM), antimycin A (50 μM) or adenine (10 mM) were added where indicated (black data points) while control plants were left untreated (grey data points). Emission in wells with Col-0 plants was averaged and subtracted from that of ATeamexpressing plants to correct for background fluorescence. Data shown and used for background subtraction is the average of 3-4 wells and error bars are SD.

**Figure 5 - figure supplement 2:**
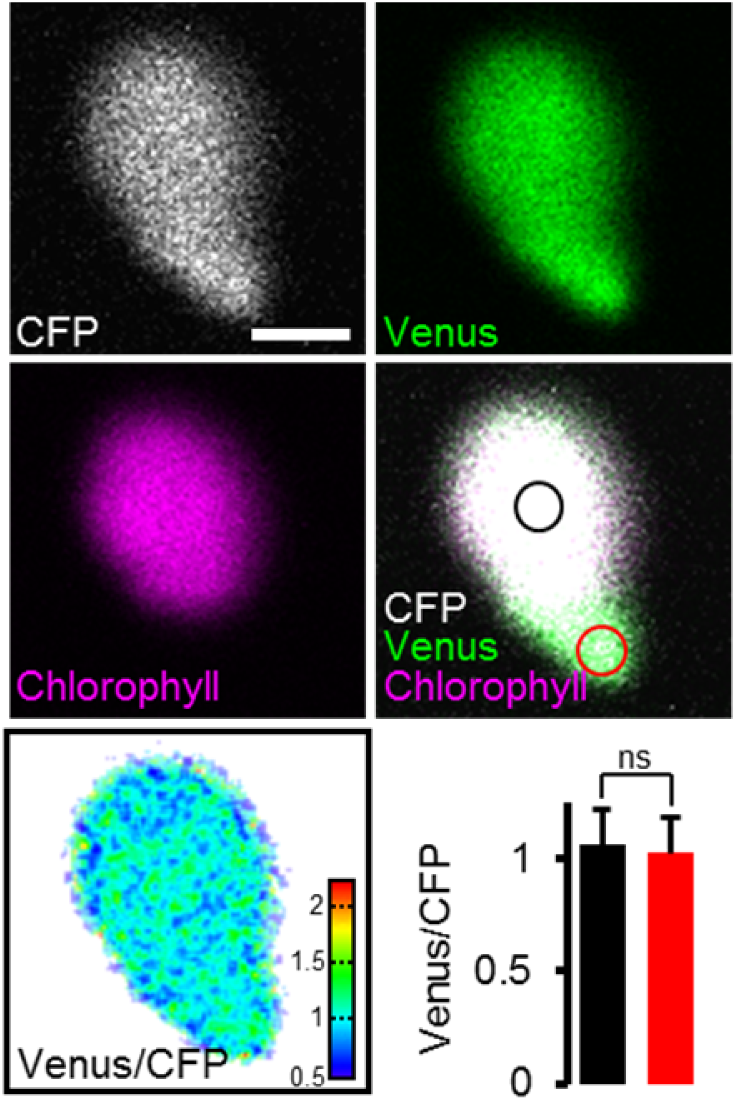
Impact of chlorophyll fluorescence on the ATeam signal. Hypocotyl of 5-day-old Arabidopsis seedlings expressing plastidic ATeam-TkTp imaged by CLSM. Venus signal, green; chlorophyll fluorescence, magenta; CFP signal, white; Venus/CFP ratio, ratiometric false-color scale. Regions of interest were defined for chloroplast centers where chlorophyll signal was present (black circle) and stromules where no chlorophyll signal was detected (red circle). The bar graph shows Venus/CFP ratios for chloroplast centers (black) and stromules (red). *n* = 11 plastids from three individual plants; error bars = SD. ns: *p* > 0.5 (*t* test). Scale bar = 2 μm.

**Figure 5 - figure supplement 3:**
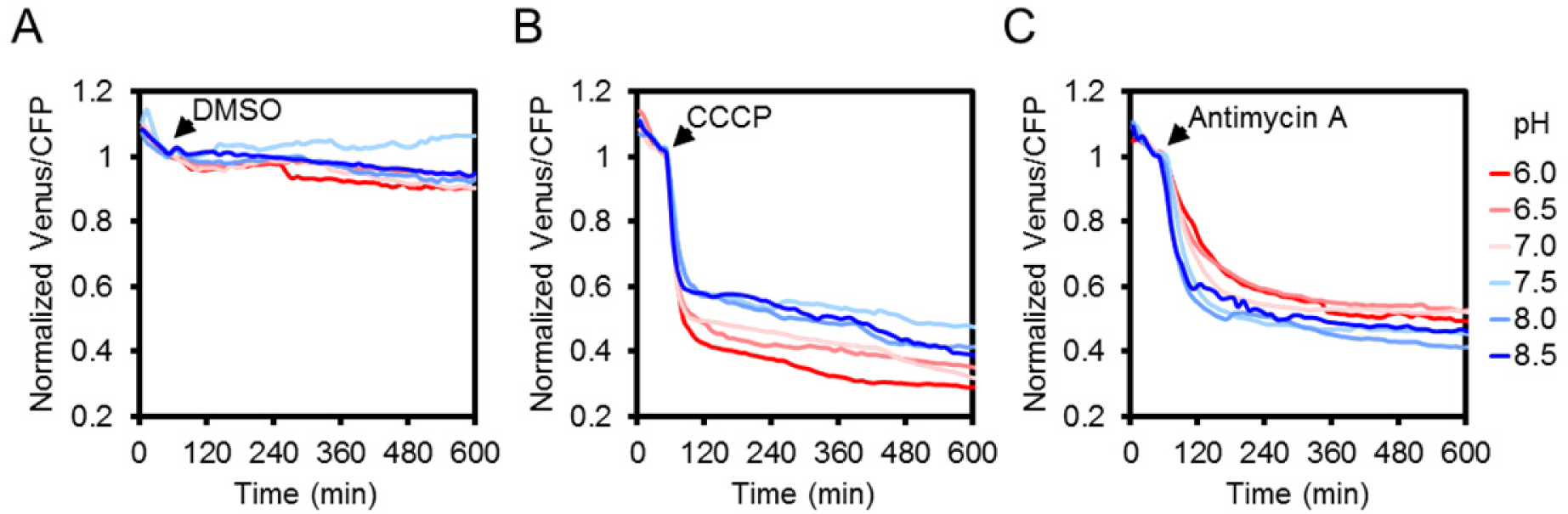
The effect of CCCP and antimycin A treatments on the ATeam response at different medium pH. Two 7-day-old Arabidopsis seedlings per well, expressing no sensor (Col-0) or cytosolic ATeam were excited at 435 ± 10 nm and the emission at 483 ± 9 nm (mseCFP) and 539 ± 6.5 nm (cp173-Venus) was recorded in imaging medium at pH values between 6.0 (red) and 8.5 (blue). DMSO as a control (A), 100 μM CCCP (B) or 100 μM antimycin A (C) were added at indicated time points. Emission in wells with Col-0 plants was averaged and subtracted from that of ATeam-expressing plants to correct for background fluorescence. Data shown and used for background subtraction are the mean of three wells.

**Figure 6 - figure supplement 1:**
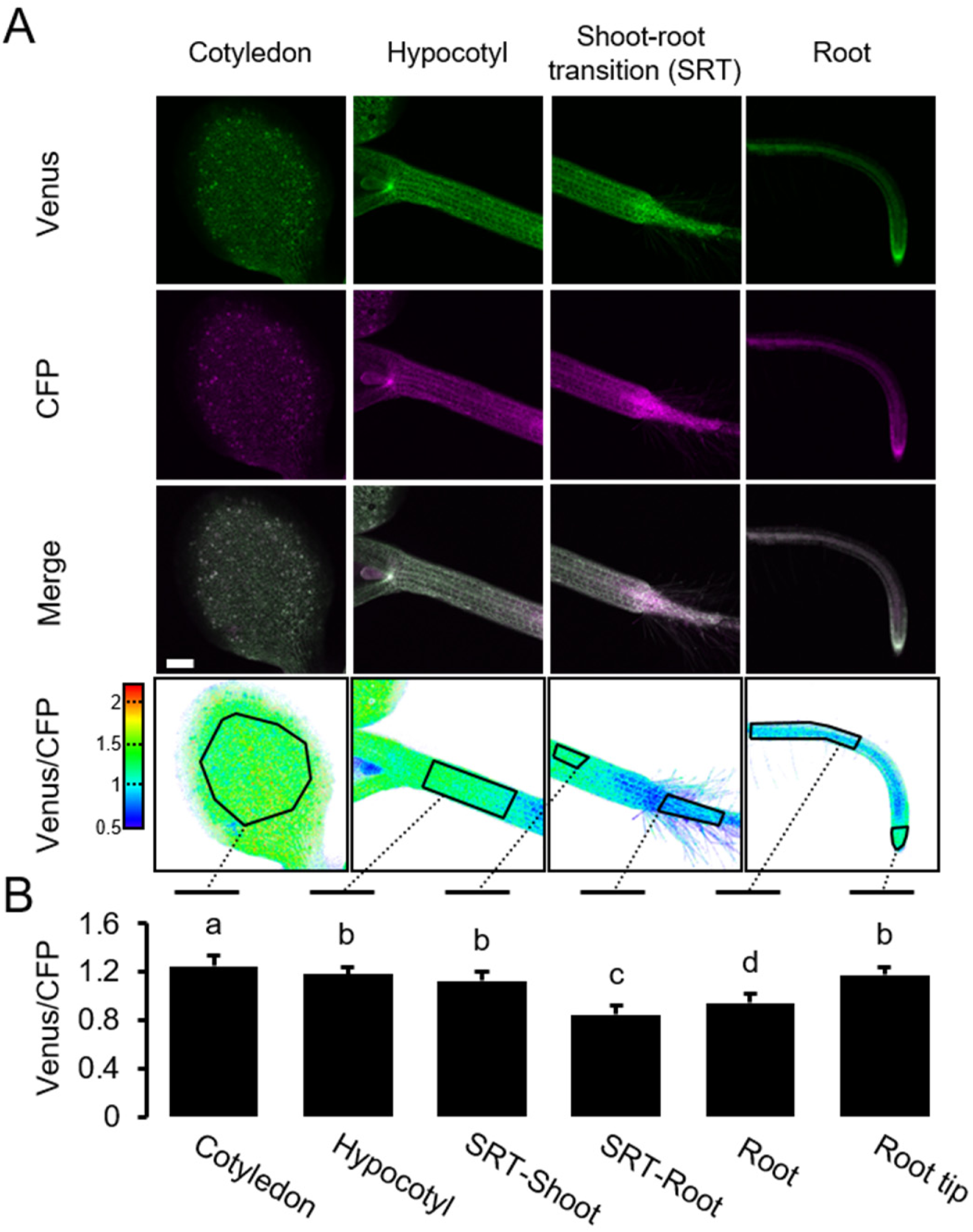
A MgATP^2−^ map of the Arabidopsis seedling. Independent repetition of experiments shown in Figure 6A,B. (A) 5-day-old Arabidopsis seedlings expressing cytosolic ATeam (line #1.1) were analysed by CLSM. Fluorescence of Venus (green) and CFP (magenta) was recorded and the ratio is plotted as a false-color image where high Venus/CFP values (red) correspond to high ATP levels. Venus/CFP ratios were analysed in the indicated regions of interest. Scale bar = 200 μm. (B) Graphs represent data from 24 individual plants; error bars = SD. Different letters indicate significant differences in a one-way ANOVA followed by the Tukey test (*p* ≤ 0.05).

**Figure 6 - figure supplement 2:**
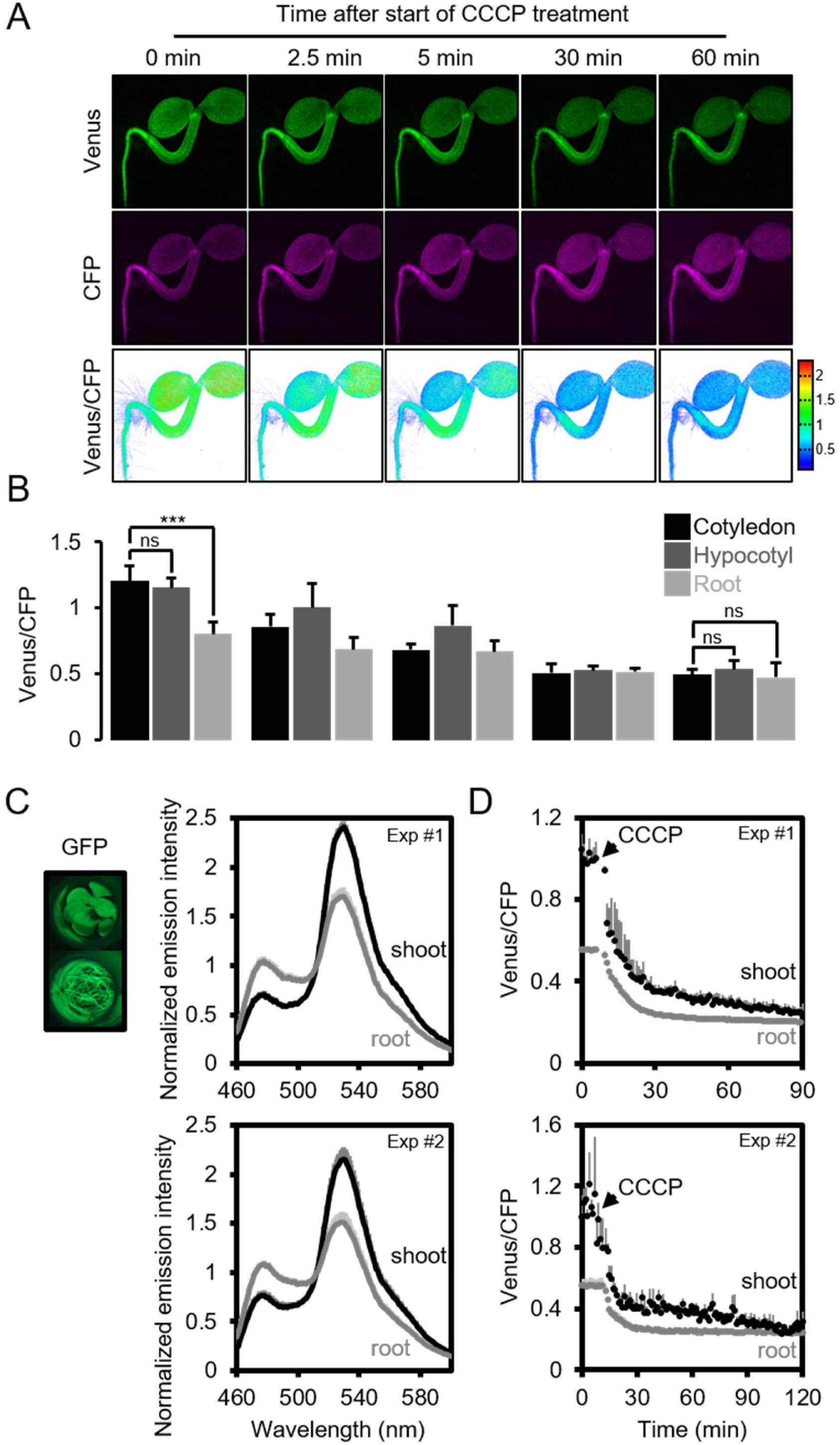
The effect of CCCP on the ATeam response in different Arabidopsis seedling tissues. (A) Five-day-old Arabidopsis seedlings expressing cytosolic ATeam were immersed in 100 μM CCCP at pH 7.5 and analysed by CLSM. Fluorescence of Venus (green) and CFP (magenta) was recorded and the ratio is plotted as a false-color image where high Venus/CFP values (red) correspond to high MgATP^2−^ levels. (B) Venus/CFP ratios at each time point after CCCP treatment were quantified in the cotyledons, hypocotyl and the upper root. *n* = 3 individual seedlings; error bars = SD. ns: *p* > 0.05, \*\*\**p* ≤ 0.001 (two-way ANOVA followed by the Tukey test). (C) Ten-day-old seedlings expressing no sensor (Col-0) or cytosolic ATeam were cut with a scalpel to separate shoots and roots that were individually submerged in imaging medium on 96-well microtiter plates. Fluorescence of plant material was checked with an epifluorescence microscope equipped with a GFP filter. Fluorescence emission spectra between 460 and 600 nm were recorded using a plate reader and an excitation wavelength of 435 ± 10 nm. The graphs show spectra of plant parts expressing cytosolic ATeam after subtraction of Col-0 fluorescence in two independent experiments (Exp #1/2). *n* = 5 wells each; error bars = SD. (D) Shoots (pooled from three seedlings per well) and roots (pooled from 10-15 seedlings per well) were excited at 435 ± 10 nm and the emission at 483 ± 9 nm (mseCFP) and 539 ± 6.5 nm (cp173-Venus) was recorded. CCCP at 100 μM was added at the indicated time point and data was plotted after subtracting fluorescence of Col-0 plant parts that were equally treated. *n* = 5 and 3 wells per tissue type in independent experiment Exp #1 and Exp #2, respectively; error bars = SD.

**Figure 10 - figure supplement 1:**
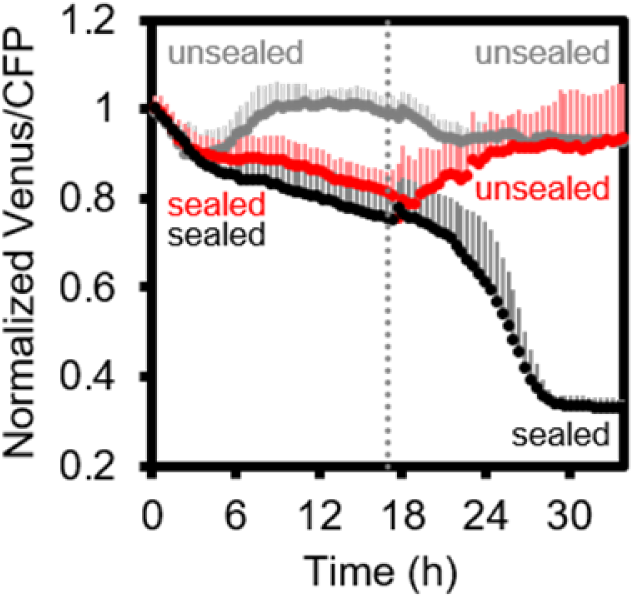
Arabidopsis MgATP^2−^ dynamics under low oxygen. Independent repetition of experiments shown in Figure 10 but using 7-day-old Arabidopsis seedlings, grown in hydroponic culture. Seedlings were submerged in imaging medium on 96-well microtiter plates. Per well, two seedlings expressing no sensor (Col-0) or cytosolic ATeam were excited at 435 ± 10 nm and the emission at 483 ± 9 nm (mseCFP) and 539 ± 6.5 nm (cp173-Venus) was recorded. Wells were either left open (grey), sealed with an oxygen-proof, transparent qPCR film (black) or sealed for 17.5 h before the film was removed to reoxygenate the samples (red). Emission in wells with Col-0 plants was averaged and subtracted from that of ATeam-expressing plants to correct for background fluorescence. Data shown and used for background subtraction is the mean of 6-10 wells and error bars are SD.

**Supplementary File 1.**
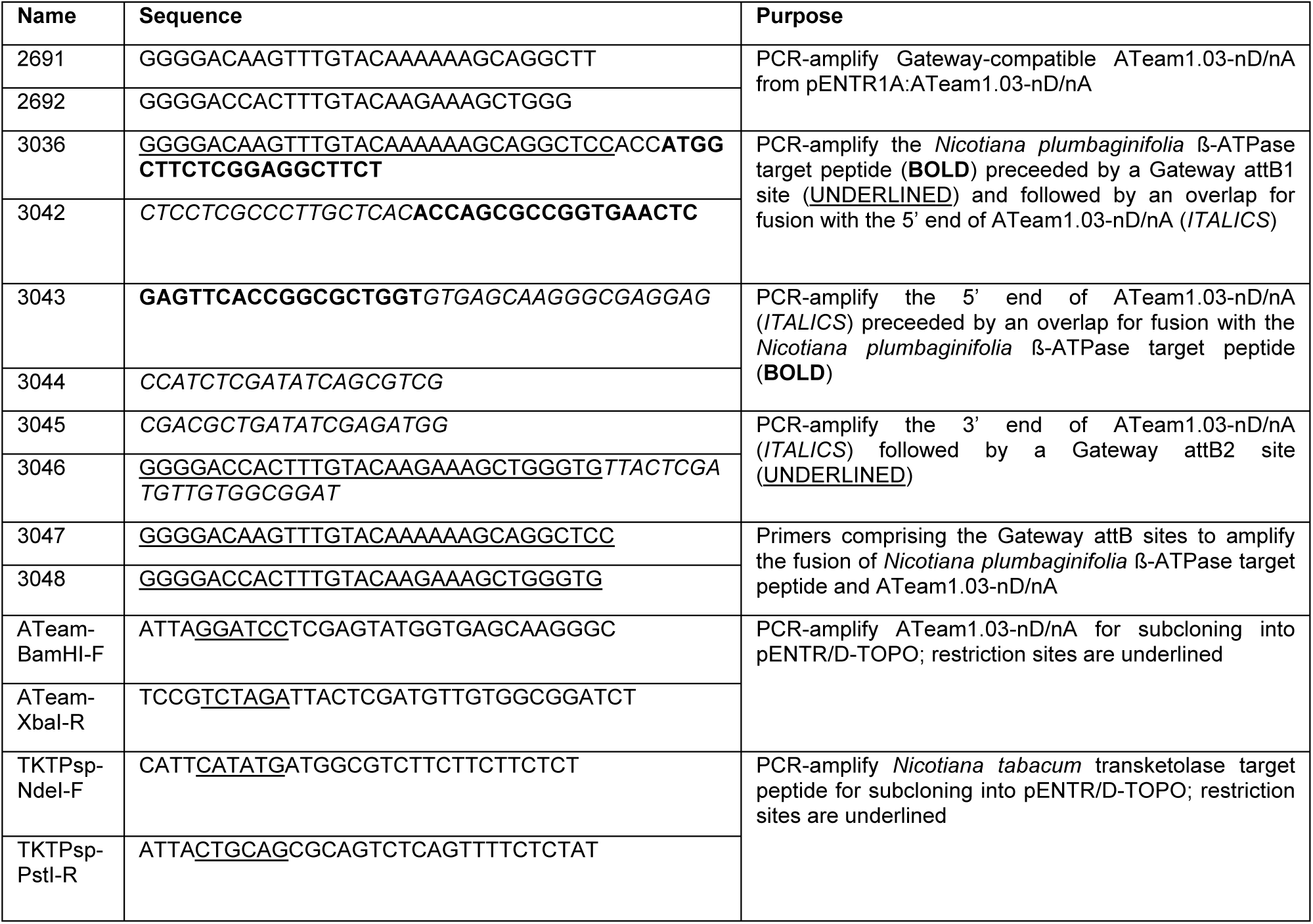

## References

Adolfsen, R., and Moudrianakis, E.N. (1978). Control of complex metal ion equilibria in biochemical reaction systems. Intrinsic and apparent stability constants of metal-adenine nucleotide complexes. J Biol Chem 253, 4378–4379.

Allen, G.J., Kwak, J.M., Chu, S.P., Llopis, J., Tsien, R.Y., Harper, J.F., and Schroeder, J.I. (1999). Cameleon calcium indicator reports cytoplasmic calcium dynamics in Arabidopsis guard cells. Plant J 19, 735–747.

Ando, T., Imamura, H., Suzuki, R., Aizaki, H., Watanabe, T., Wakita, T., and Suzuki, T. (2012). Visualization and measurement of ATP levels in living cells replicating hepatitis C virus genome RNA. PLoS Pathog 8, e1002561.

Attucci, S., Carde, J.P., Raymond, P., Saint-Ges, V., Spiteri, A., and Pradet, A. (1991). Oxidative phosphorylation by mitochondria extracted from dry sunflower seeds. Plant Physiol 95, 390–398.

Bailleul, B., Berne, N., Murik, O., Petroutsos, D., Prihoda, J., Tanaka, A., Villanova, V., Bligny, R., Flori, S., Falconet, D., Krieger-Liszkay, A., Santabarbara, S., Rappaport, F., Joliot, P., Tirichine, L., Falkowski, P.G., Cardol, P., Bowler, C., and Finazzi, G. (2015). Energetic coupling between plastids and mitochondria drives CO2 assimilation in diatoms. Nature 524, 366–369.

Behera, S., Wang, N., Zhang, C., Schmitz-Thom, I., Strohkamp, S., Schultke, S., Hashimoto, K., Xiong, L., and Kudla, J. (2015). Analyses of Ca^2+^ dynamics using a ubiquitin-10 promoter-driven Yellow Cameleon 3.6 indicator reveal reliable transgene expression and differences in cytoplasmic Ca^2+^ responses in Arabidopsis and rice (*Oryza sativa*) roots. New Phytol 206, 751–760.

Berg, J., Hung, Y.P., and Yellen, G. (2009). A genetically encoded fluorescent reporter of ATP:ADP ratio. Nature Methods 6, 161–166.

Bertani, G. (1951). Studies on lysogenesis. I. The mode of phage liberation by lysogenic *Escherichia coli*. J Bacteriol 62, 293–300.

Bidel, L.P., Renault, P., Pages, L., and Riviere, L.M. (2000). Mapping meristem respiration of *Prunus persica* (L.) Batsch seedlings: potential respiration of the meristems, O2 diffusional constraints and combined effects on root growth. J Exp Bot 51, 755–768.

Birk, J., Ramming, T., Odermatt, A., and Appenzeller-Herzog, C. (2013). Green fluorescent protein-based monitoring of endoplasmic reticulum redox poise. Front Genet 4, 108.

Birkenhead, K., Walker, D., and Foyer, C. (1982). The intracellular distribution of adenylate kinase in the leaves of spinach, wheat and barley. Planta 156, 171–175.

Bradford, M.M. (1976). A rapid and sensitive method for the quantitation of microgram quantities of protein utilizing the principle of protein-dye binding. Anal Biochem 72, 248–254.

Brunoud, G., Wells, D.M., Oliva, M., Larrieu, A., Mirabet, V., Burrow, A.H., Beeckman, T., Kepinski, S., Traas, J., Bennett, M.J., and Vernoux, T. (2012). A novel sensor to map auxin response and distribution at high spatio-temporal resolution. Nature 482, 103–106.

Busch, K., and Ninnemann, H. (1996). Characterization of adenylate kinase in intact mitochondria of fertile and male sterile potato (*Solanum tuberosum* L.). Plant Science 116, 1–8.

Candeo, A., Doccula, F.G., Valentini, G., Bassi, A., and Costa, A. (2017). Light sheet fluorescence microscopy quantifies calcium oscillations in root hairs of *Arabidopsis thaliana*. Plant Cell Physiol published online ahead of print: doi:10.1093/pcp/pcx045.

Chaudhuri, B., Hormann, F., Lalonde, S., Brady, S.M., Orlando, D.A., Benfey, P., and Frommer, W.B. (2008). Protonophore- and pH-insensitive glucose and sucrose accumulation detected by FRET nanosensors in Arabidopsis root tips. Plant J 56, 948–962.

Choi, J., Tanaka, K., Cao, Y., Qi, Y., Qiu, J., Liang, Y., Lee, S.Y., and Stacey, G. (2014a). Identification of a plant receptor for extracellular ATP. Science 343, 290–294.

Choi, W.G., Toyota, M., Kim, S.H., Hilleary, R., and Gilroy, S. (2014b). Salt stress-induced Ca^2+^ waves are associated with rapid, long-distance root-to-shoot signaling in plants. Proc Natl Acad Sci U S A 111, 6497–6502.

Clough, S.J., and Bent, A.F. (1998). Floral dip: a simplified method for Agrobacterium-mediated transformation of *Arabidopsis thaliana*. Plant J 16, 735–743.

Considine, M.J., Diaz-Vivancos, P., Kerchev, P., Signorelli, S., Agudelo-Romero, P., Gibbs, D.J., and Foyer, C.H. (2017). Learning to breathe: developmental phase transitions in oxygen status. Trends Plant Sci 22, 140–153.

Costa, A., Candeo, A., Fieramonti, L., Valentini, G., and Bassi, A. (2013). Calcium dynamics in root cells of *Arabidopsis thaliana* visualized with selective plane illumination microscopy. PLoS One 8, e75646.

Day, D.A., Arron, G.P., and Laties, G.G. (1979). Enzyme distribution in potato mitochondria. Journal of Experimental Botany 30, 539–549.

De Michele, R., Carimi, F., and Frommer, W.B. (2014). Mitochondrial biosensors. Int J Biochem Cell Biol 48, 39–44.

Dejonghe, W., and Russinova, E. (2017). Plant chemical genetics: from phenotype-based screens to synthetic biology. Plant Physiol e-published ahead of print, doi:10.1104/pp.1116.01805.

Deuschle, K., Chaudhuri, B., Okumoto, S., Lager, I., Lalonde, S., and Frommer, W.B. (2006). Rapid metabolism of glucose detected with FRET glucose nanosensors in epidermal cells and intact roots of Arabidopsis RNA-silencing mutants. Plant Cell 18, 2314–2325.

Dickinson, D.B. (1965). Germination of lily pollen: respiration and tube growth. Science 150, 1818–1819.

Earley, K.W., Haag, J.R., Pontes, O., Opper, K., Juehne, T., Song, K., and Pikaard, C.S. (2006). Gateway-compatible vectors for plant functional genomics and proteomics. Plant J 45, 616–629.

Flügge, U.I. (1998). Metabolite transporters in plastids. Curr Opin Plant Biol 1, 201–206.

Fricker, M.D. (2016). Quantitative redox imaging software. Antioxid Redox Signal 24, 752–762.

Gardeström, P., and Wigge, B. (1988). Influence of photorespiration on ATP/ADP ratios in the chloroplasts, mitochondria, and cytosol, studied by rapid fractionation of barley (*Hordeum vulgare*) protoplasts. Plant Physiol 88, 69–76.

Gardeström, P., and Igamberdiev, A.U. (2016). The origin of cytosolic ATP in photosynthetic cells. Physiol Plant 157, 367–379.

Geigenberger, P. (2003). Response of plant metabolism to too little oxygen. Curr Opin Plant Biol 6, 247–256.

Geigenberger, P., Fernie, A.R., Gibon, Y., Christ, M., and Stitt, M. (2000). Metabolic activity decreases as an adaptive response to low internal oxygen in growing potato tubers. Biol Chem 381, 723–740.

Gout, E., Rebeille, F., Douce, R., and Bligny, R. (2014). Interplay of Mg^2+^, ADP, and ATP in the cytosol and mitochondria: unravelling the role of Mg^2+^ in cell respiration. Proc Natl Acad Sci U S A 111, E4560–4567.

Gout, E., Boisson, A., Aubert, S., Douce, R., and Bligny, R. (2001). Origin of the cytoplasmic pH changes during anaerobic stress in higher plant cells. Carbon-13 and phosphorous-31 nuclear magnetic resonance studies. Plant Physiol 125, 912–925.

Haferkamp, I., Fernie, A.R., and Neuhaus, H.E. (2011). Adenine nucleotide transport in plants: much more than a mitochondrial issue. Trends Plant Sci 16, 507–515.

Hatsugai, N., Perez Koldenkova, V., Imamura, H., Noji, H., and Nagai, T. (2012). Changes in cytosolic ATP levels and intracellular morphology during bacteria-induced hypersensitive cell death as revealed by real-time fluorescence microscopy imaging. Plant Cell Physiol 53, 1768–1775.

Hepler, P.K., Vidali, L., and Cheung, A.Y. (2001). Polarized cell growth in higher plants. Annu Rev Cell Dev Biol 17, 159–187.

Igamberdiev, A.U., and Kleczkowski, L.A. (2003). Membrane potential, adenylate levels and Mg^2+^ are interconnected via adenylate kinase equilibrium in plant cells. Biochim Biophys Acta 1607, 111–119.

Igamberdiev, A.U., Bykova, N.V., Lea, P.J., and Gardeström, P. (2001). The role of photo respiration in redox and energy balance of photosynthetic plant cells: A study with a barley mutant deficient in glycine decarboxylase. Physiol Plant 111, 427–438.

Imamura, H., Nhat, K.P., Togawa, H., Saito, K., Iino, R., Kato-Yamada, Y., Nagai, T., and Noji, H. (2009). Visualization of ATP levels inside single living cells with fluorescence resonance energy transfer-based genetically encoded indicators. Proc Natl Acad Sci U S A 106, 15651–15656.

Jones, A.M., Danielson, J.A., Manojkumar, S.N., Lanquar, V., Grossmann, G., and Frommer, W.B. (2014). Abscisic acid dynamics in roots detected with genetically encoded FRET sensors. Elife 3, e01741.

Karimi, M., Inze, D., and Depicker, A. (2002). GATEWAY vectors for Agrobacterium-mediated plant transformation. Trends Plant Sci 7, 193–195.

Keinath, N.F., Waadt, R., Brugman, R., Schroeder, J.I., Grossmann, G., Schumacher, K., and Krebs, M. (2015). Live cell imaging with R-GECO1 sheds light on flg22- and chitin-induced transient [Ca^2+^]_cyt_ patterns in *Arabidopsis*. Mol Plant 8, 1188–1200.

Khlyntseva, S.V., Bazel’, Y.R., Vishnikin, A.B., and Andruch, V. (2009). Methods for the determination of adenosine triphosphate and other adenine nucleotides. Journal of Analytical Chemistry 64, 657–673.

Kim, S.Y., Sivaguru, M., and Stacey, G. (2006). Extracellular ATP in plants. Visualization, localization, and analysis of physiological significance in growth and signaling. Plant Physiol 142, 984–992.

Kotera, I., Iwasaki, T., Imamura, H., Noji, H., and Nagai, T. (2010). Reversible dimerization of *Aequorea victoria* fluorescent proteins increases the dynamic range of FRET-based indicators. ACS Chem Biol 5, 215–222.

Krebs, M., Held, K., Binder, A., Hashimoto, K., Den Herder, G., Parniske, M., Kudla, J., and Schumacher, K. (2012). FRET-based genetically encoded sensors allow high-resolution live cell imaging of Ca^2+^ dynamics. Plant J 69, 181–192.

Krömer, S., and Heldt, H.W. (1991). On the role of mitochondrial oxidative phosphorylation in photosynthesis metabolism as studied by the effect of oligomycin on photosynthesis in protoplasts and leaves of barley (*Hordeum vulgare*). Plant Physiol 95, 1270–1276.

Krömer, S., Malmberg, G., and Gardeström, P. (1993). Mitochondrial contribution to photosynthetic metabolism (A study with barley (*Hordeum vulgare* L.) leaf protoplasts at different light intensities and CO2 concentrations). Plant Physiol 102, 947–955.

Li, J., Yu, Q., Ahooghalandari, P., Gribble, F.M., Reimann, F., Tengholm, A., and Gylfe, E. (2015). Submembrane ATP and Ca^2+^ kinetics in alpha-cells: unexpected signaling for glucagon secretion. FASEB J 29, 3379–3388.

Li, S., Yamada, M., Han, X., Ohler, U., and Benfey, P.N. (2016). High-resolution expression map of the Arabidopsis root reveals alternative splicing and lincRNA regulation. Dev Cell 39, 508–522.

Lilley, R.M., Stitt, M., Mader, G., and Heldt, H.W. (1982). Rapid fractionation of wheat leaf protoplasts using membrane filtration: the determination of metabolite levels in the chloroplasts, cytosol, and mitochondria. Plant Physiol 70, 965–970.

Loef, I., Stitt, M., and Geigenberger, P. (2001). Increased levels of adenine nucleotides modify the interaction between starch synthesis and respiration when adenine is supplied to discs from growing potato tubers. Planta 212, 782–791.

Logan, D.C., and Leaver, C.J. (2000). Mitochondria-targeted GFP highlights the heterogeneity of mitochondrial shape, size and movement within living plant cells. J Exp Bot 51, 865–871.

Lorenz, A., Lorenz, M., Vothknecht, U.C., Niopek-Witz, S., Neuhaus, H.E., and Haferkamp, I. (2015). *In vitro* analyses of mitochondrial ATP/phosphate carriers from *Arabidopsis thaliana* revealed unexpected Ca^2+^-effects. BMC Plant Biol 15, 238.

Loro, G., Drago, I., Pozzan, T., Schiavo, F.L., Zottini, M., and Costa, A. (2012). Targeting of Cameleons to various subcellular compartments reveals a strict cytoplasmic/mitochondrial Ca^2+^ handling relationship in plant cells. Plant J 71, 1–13.

Loro, G., Wagner, S., Doccula, F.G., Behera, S., Weinl, S., Kudla, J., Schwarzländer, M., Costa, A., and Zottini, M. (2016). Chloroplast-specific *in vivo* Ca^2+^ imaging using Yellow Cameleon fluorescent protein sensors reveals organelle-autonomous Ca^2+^ signatures in the stroma. Plant Physiol 171, 2317–2330.

Luo, Y., Scholl, S., Doering, A., Zhang, Y., Irani, N.G., Di Rubbo, S., Neumetzler, L., Krishnamoorthy, P., Van Houtte, I., Mylle, E., Bischoff, V., Vernhettes, S., Winne, J., Friml, J., Stierhof, Y.D., Schumacher, K., Persson, S., and Russinova, E. (2015). V-ATPase activity in the TGN/EE is required for exocytosis and recycling in *Arabidopsis*. Nat Plants 1, 15094.

Manfredi, G., Yang, L., Gajewski, C.D., and Mattiazzi, M. (2002). Measurements of ATP in mammalian cells. Methods 26, 317–326.

Marty, L., Siala, W., Schwarzländer, M., Fricker, M.D., Wirtz, M., Sweetlove, L.J., Meyer, Y., Meyer, A.J., Reichheld, J.P., and Hell, R. (2009). The NADPH-dependent thioredoxin system constitutes a functional backup for cytosolic glutathione reductase in *Arabidopsis*. Proc Natl Acad Sci U S A 106, 9109–9114.

Merrins, M.J., Poudel, C., McKenna, J.P., Ha, J., Sherman, A., Bertram, R., and Satin, L.S. (2016). Phase analysis of metabolic oscillations and membrane potential in pancreatic islet beta-cells. Biophys J 110, 691–699.

Meyer, A.J., Brach, T., Marty, L., Kreye, S., Rouhier, N., Jacquot, J.P., and Hell, R. (2007). Redox-sensitive GFP in *Arabidopsis thaliana* is a quantitative biosensor for the redox potential of the cellular glutathione redox buffer. Plant J 52, 973–986.

Monne, M., Miniero, D.V., Obata, T., Daddabbo, L., Palmieri, L., Vozza, A., Nicolardi, M.C., Fernie, A.R., and Palmieri, F. (2015). Functional characterization and organ distribution of three mitochondrial ATP-Mg/Pi carriers in *Arabidopsis thaliana*. Biochim Biophys Acta 1847, 1220–1230.

Morgan, B., Ezerina, D., Amoako, T.N., Riemer, J., Seedorf, M., and Dick, T.P. (2013). Multiple glutathione disulfide removal pathways mediate cytosolic redox homeostasis. Nat Chem Biol 9, 119–125.

Morgan, B., Van Laer, K., Owusu, T.N., Ezerina, D., Pastor-Flores, D., Amponsah, P.S., Tursch, A., and Dick, T.P. (2016). Real-time monitoring of basal H2O2 levels with peroxiredoxin-based probes. Nat Chem Biol 12, 437–443.

Murashige, T., and Skoog, F. (1962). A revised medium for rapid growth and bio assays with tobacco tissue cultures. Physiologia Plantarum 15, 473–497.

Mustroph, A., Boamfa, E.I., Laarhoven, L.J., Harren, F.J., Albrecht, G., and Grimm, B. (2006). Organ-specific analysis of the anaerobic primary metabolism in rice and wheat seedlings. I: Dark ethanol production is dominated by the shoots. Planta 225, 103–114.

Nakano, M., Imamura, H., Nagai, T., and Noji, H. (2011). Ca^2+^ regulation of mitochondrial ATP synthesis visualized at the single cell level. ACS Chem Biol 6, 709–715.

Neuhaus, H.E., Thom, E., Möhlmann, T., Steup, M., and Kampfenkel, K. (1997). Characterization of a novel eukaryotic ATP/ADP translocator located in the plastid envelope of *Arabidopsis thaliana* L. Plant J 11, 73–82.

O’Sullivan, W.J., and Perrin, D.D. (1961). The stability constants of MgATP^−2^ ion. Biochim Biophys Acta 52, 612–614.

Palmer, A.E., and Tsien, R.Y. (2006). Measuring calcium signaling using genetically targetable fluorescent indicators. Nat Protoc 1, 1057–1065.

Patel, A., Malinovska, L., Saha, S., Wang, J., Alberti, S., Krishnan, Y., and Hyman, A.A. (2017). ATP as a biological hydrotrope. Science 356, 753–756.

Pei, Z.M., Murata, Y., Benning, G., Thomine, S., Klusener, B., Allen, G.J., Grill, E., and Schroeder, J.I. (2000). Calcium channels activated by hydrogen peroxide mediate abscisic acid signalling in guard cells. Nature 406, 731–734.

Pradet, A., and Raymond, P. (1983). Adenine nucleotide ratios and adenylate energy charge in energy metabolism. Annual Review of Plant Physiology 34, 199–224.

Ratcliffe, R.G. (1997). *In vivo* NMR studies of the metabolic response of plant tissues to anoxia. Annals of Botany 79, 39–48.

Reiser, J., Linka, N., Lemke, L., Jeblick, W., and Neuhaus, H.E. (2004). Molecular physiological analysis of the two plastidic ATP/ADP transporters from Arabidopsis. Plant Physiol 136, 3524–3536.

Rich, P. (2003). Chemiosmotic coupling: The cost of living. Nature 421, 583.

Roberts, J., Aubert, S., Gout, E., Bligny, R., and Douce, R. (1997). Cooperation and competition between adenylate kinase, nucleoside diphosphokinase, electron transport, and ATP synthase in plant mitochondria studied by ^31^P-nuclear magnetic resonance. Plant Physiol 113, 191–199.

Rosenwasser, S., Rot, I., Meyer, A.J., Feldman, L., Jiang, K., and Friedman, H. (2010). A fluorometer-based method for monitoring oxidation of redox-sensitive GFP (roGFP) during development and extended dark stress. Physiol Plant 138, 493–502.

Rosenwasser, S., Rot, I., Sollner, E., Meyer, A.J., Smith, Y., Leviatan, N., Fluhr, R., and Friedman, H. (2011). Organelles contribute differentially to reactive oxygen species-related events during extended darkness. Plant Physiol 156, 185–201.

Schulte, A., Lorenzen, I., Böttcher, M., and Plieth, C. (2006). A novel fluorescent pH probe for expression in plants. Plant Methods 2, 7.

Schulz, G.E. (1987). Structural and functional relationships in the adenylate kinase family. Cold Spring Harb Symp Quant Biol 52, 429–439.

Schwarzländer, M., Fricker, M.D., and Sweetlove, L.J. (2009). Monitoring the *in vivo* redox state of plant mitochondria: effect of respiratory inhibitors, abiotic stress and assessment of recovery from oxidative challenge. Biochim Biophys Acta 1787, 468–475.

Schwarzländer, M., Logan, D.C., Fricker, M.D., and Sweetlove, L.J. (2011). The circularly permuted yellow fluorescent protein cpYFP that has been used as a superoxide probe is highly responsive to pH but not superoxide in mitochondria: implications for the existence of superoxide ‘flashes’. Biochem J 437, 381–387.

Schwarzländer, M., Dick, T.P., Meyer, A.J., and Morgan, B. (2016). Dissecting redox biology using fluorescent protein sensors. Antioxid Redox Signal 24, 680–712.

Schwarzländer, M., Logan, D.C., Johnston, I.G., Jones, N.S., Meyer, A.J., Fricker, M.D., and Sweetlove, L.J. (2012). Pulsing of membrane potential in individual mitochondria: a stress-induced mechanism to regulate respiratory bioenergetics in *Arabidopsis*. Plant Cell 24, 1188–1201.

Schwarzländer, M., Fricker, M.D., Müller, C., Marty, L., Brach, T., Novak, J., Sweetlove, L.J., Hell, R., and Meyer, A.J. (2008). Confocal imaging of glutathione redox potential in living plant cells. J Microsc 231, 299–316.

Stitt, M., Lilley, R.M., and Heldt, H.W. (1982). Adenine nucleotide levels in the cytosol, chloroplasts, and mitochondria of wheat leaf protoplasts. Plant Physiol 70, 971–977.

Storer, A.C., and Cornish-Bowden, A. (1976). Concentration of MgATP^2−^ and other ions in solution. Calculation of the true concentrations of species present in mixtures of associating ions. Biochem J 159, 1–5.

Sweetlove, L.J., Taylor, N.L., and Leaver, C.J. (2007). Isolation of intact, functional mitochondria from the model plant *Arabidopsis thaliana*. Methods Mol Biol 372, 125–136.

Sweetlove, L.J., Nielsen, J., and Fernie, A.R. (2017). Engineering central metabolism - a grand challenge for plant biologists. Plant J.

Sweetlove, L.J., Heazlewood, J.L., Herald, V., Holtzapffel, R., Day, D.A., Leaver, C.J., and Millar, A.H. (2002). The impact of oxidative stress on Arabidopsis mitochondria. Plant J 32, 891–904.

Taiz, L., Zeiger, E., Møller, I.M., and Murphy, A. (2015). Plant physiology and development, 6th edition. (Sunderland, CT: Sinauer Associates).

Tanaka, K., Gilroy, S., Jones, A.M., and Stacey, G. (2010). Extracellular ATP signaling in plants. Trends Cell Biol 20, 601–608.

Tantama, M., Martinez-Francois, J.R., Mongeon, R., and Yellen, G. (2013). Imaging energy status in live cells with a fluorescent biosensor of the intracellular ATP-to-ADP ratio. Nat Commun 4, 2550.

Tarasov, A.I., Semplici, F., Ravier, M.A., Bellomo, E.A., Pullen, T.J., Gilon, P., Sekler, I., Rizzuto, R., and Rutter, G.A. (2012). The mitochondrial Ca^2+^ uniporter MCU is essential for glucose-induced ATP increases in pancreatic beta-cells. PLoS One 7, e39722.

Uslu, V.V., and Grossmann, G. (2016). The biosensor toolbox for plant developmental biology. Curr Opin Plant Biol 29, 138–147.

van Dongen, J.T., and Licausi, F. (2015). Oxygen sensing and signaling. Annu Rev Plant Biol 66, 345–367.

van Dongen, J.T., Schurr, U., Pfister, M., and Geigenberger, P. (2003). Phloem metabolism and function have to cope with low internal oxygen. Plant Physiol 131, 1529–1543.

van Dongen, J.T., Frohlich, A., Ramirez-Aguilar, S.J., Schauer, N., Fernie, A.R., Erban, A., Kopka, J., Clark, J., Langer, A., and Geigenberger, P. (2009). Transcript and metabolite profiling of the adaptive response to mild decreases in oxygen concentration in the roots of arabidopsis plants. Ann Bot 103, 269–280.

Waadt, R., Hitomi, K., Nishimura, N., Hitomi, C., Adams, S.R., Getzoff, E.D., and Schroeder, J.I. (2014). FRET-based reporters for the direct visualization of abscisic acid concentration changes and distribution in Arabidopsis. Elife 3, e01739.

Wagner, S., Nietzel, T., Aller, I., Costa, A., Fricker, M.D., Meyer, A.J., and Schwarzländer, M. (2015a). Analysis of plant mitochondrial function using fluorescent protein sensors. Methods Mol Biol 1305, 241–252.

Wagner, S., Behera, S., De Bortoli, S., Logan, D.C., Fuchs, P., Carraretto, L., Teardo, E., Cendron, L., Nietzel, T., Füssl, M., Doccula, F.G., Navazio, L., Fricker, M.D., Van Aken, O., Finkemeier, I., Meyer, A.J., Szabo, I., Costa, A., and Schwarzländer, M. (2015b). The EF-hand Ca^2+^ binding protein MICU choreographs mitochondrial Ca^2+^ dynamics in Arabidopsis. Plant Cell 27, 3190–3212.

Wend, S., Dal Bosco, C., Kampf, M.M., Ren, F., Palme, K., Weber, W., Dovzhenko, A., and Zurbriggen, M.D. (2013). A quantitative ratiometric sensor for time-resolved analysis of auxin dynamics. Sci Rep 3, 2052.

Wirtz, M., and Hell, R. (2003). Production of cysteine for bacterial and plant biotechnology: application of cysteine feedback-insensitive isoforms of serine acetyltransferase. Amino Acids 24, 195–203.

Xia, J.H., and Saglio, P.H. (1992). Lactic acid efflux as a mechanism of hypoxic acclimation of maize root tips to anoxia. Plant Physiol 100, 40–46.

Yagi, H., Kajiwara, N., Tanaka, H., Tsukihara, T., Kato-Yamada, Y., Yoshida, M., and Akutsu, H. (2007). Structures of the thermophilic F1-ATPase ε subunit suggesting ATP-regulated arm motion of its C-terminal domain in F1. Proc Natl Acad Sci U S A 104, 11233–11238.

Yang, H., Bogner, M., Stierhof, Y.D., and Ludewig, U. (2010). H^+^-independent glutamine transport in plant root tips. PLoS One 5, e8917.

Yoshida, T., Kakizuka, A., and Imamura, H. (2016). BTeam, a novel BRET-based biosensor for the accurate quantification of ATP concentration within living cells. Sci Rep 6, 39618.

Zabalza, A., van Dongen, J.T., Froehlich, A., Oliver, S.N., Faix, B., Gupta, K.J., Schmalzlin, E., Igal, M., Orcaray, L., Royuela, M., and Geigenberger, P. (2009). Regulation of respiration and fermentation to control the plant internal oxygen concentration. Plant Physiol 149, 1087–1098.

Zancani, M., Casolo, V., Vianello, A., and Macrȶ, F. (2001). Involvement of apyrase in the regulation of the adenylate pool by adenylate kinase in plant mitochondria. Plant Science 161, 927–933.

